# A new group of LysM-RLKs involved in symbiotic signal perception and arbuscular mycorrhiza establishment

**DOI:** 10.1101/2024.03.06.583654

**Authors:** Yi Ding, Virginie Gasciolli, Lauréna Medioni, Mégane Gaston, Annélie de-Regibus, Céline Rem-blière, Jean Jacques Bono, Julie Cullimore, Marion Dalmais, Christine Saffray, Solène Mazeau, Abdelhafid Bendahmane, Richard Sibout, Michiel Vandenbussche, Jacques Rouster, Tongming Wang, Guanghua He, Arnaud Masselin, Sylvain Cottaz, Sébastien Fort, Benoit Lefebvre

## Abstract

Lipo-chitooligosaccharides (LCO) and short-chain chitooligosaccharides (CO) are produced by arbuscular mycorrhizal fungi (AMF) and activate the plant symbiosis signalling pathway, which is essential for mycorrhiza formation. High affinity LCO receptors belonging to the LysM receptor-like kinase (LysM-RLK) phylogenetic group *LYR-IA* play a role in AM establishment, but no plant high affinity short-chain CO receptors have yet been identified. Here we studied members of the uncharacterized *LYR-IB* group, and found that they show high affinity for LCO, short- and long-chain CO, and play a complementary role with the *LYR-IA* LCO receptors for AM establishment. While *LYR-IB* knock out mutants had a reduced AMF colonization in several species, constitutive/ectopic expression in wheat increased AMF colonization. *LYR-IB* function is conserved in all tested angiosperms, but in most japonica rice a deletion creates a frameshift in the gene, explaining differences in AM phenotypes between rice and other monocot single *LYR-IA* mutants. In conclusion, we identified a class of LysM-RLK receptors in angiosperms with new biochemical properties and a role in both LCO and CO perception for AM establishment.

## Introduction

Arbuscular mycorrhiza (AM) is an ancient mutualistic symbiosis between Glomeromycotina fungi and most plant species, including crops such as rice and wheat. Arbuscular mycorrhizal fungi (AMF) provide plants with nutrients gathered in the soil through an extended extraradical mycelium in exchange for carbohydrates and lipids^1^. Host root colonization by AMF relies on a signalling pathway called the common symbiosis signalling pathway (CSSP)^2^. Two types of chitinic signal molecules produced by AMF, lipo-chitooligosaccharides (LCO)^3^ and short-chain chitooligosaccharides (CO)^4^ induce at nM concentrations oscillations in calcium concentration in and around root cell nuclei (calcium spiking)^5^, a hallmark of CSSP activation, and they stimulate AMF colonization when added exogenously^3,6^.

During evolution, the CSSP was recruited for nodule organogenesis and colonization by nitrogen-fixing bacteria in nodulating plant species^2^. In most legumes, including *Medicago truncatula*, the CSSP is activated by specific LCO structures produced by their nitrogen-fixing rhizobial partner leading to calcium spiking and nodulation.

Both LCO and short-chain CO are composed of 4 to 5 N-acetylglucosamine (GlcNAc) residues. In addition, LCO contain a lipid on the terminal non-reducing sugar. In contrast to LCO and short-chain CO, longer CO (mainly those containing 7 or 8 GlcNAc) and peptidoglycan (PGN) fragments (a polymer of N-acetylmuramic acid and GlcNAc), induce defence responses in plants.

Although inducing different types of responses, all these GlcNAc-containing molecules are perceived by plant specific plasma membrane localized proteins containing three Lysin motifs (LysM) in their extracellular region (ECR). LysM receptor-like kinases (LysM-RLKs) also have a transmembrane domain (TM) and an intracellular region (ICR) with an active kinase domain (LYK type) or an inactive kinase domain (LYR type), while LysM-receptor like proteins (LysM-RLPs) have a GPI anchor site and no ICR (LYM type). LYR and LYM proteins interact with LYK proteins to form receptor complexes in which in most cases the LYR or the LYM has high affinity and specificity for a ligand and the LYK lower affinity and specificity for ligands, but a critical role in signal transduction^7^. Conversely, LYR and LYM are generally involved in specific signalling pathways, while, through interaction with several LYR and LYM, LYK are involved in multiple signalling pathways leading to symbiosis or defence^7^. LysM-RLK/P numbers vary between species (from 8 to more than 20) but several phylogenetic groups are conserved between angiosperms^7^.

Several LysM-RLK/Ps have been described as high affinity LCO or long-chain CO binding proteins^8–17^. Overall, the ligand binding properties of the proteins are conserved between orthologs belonging to conserved phylogenetic groups in angiosperms^7^, except some exceptions likely resulting from evolutionary processes associated with gain or loss of symbiosis abilities^2,17^. For example, members of the *LYM-II* group have high affinity for long-chain CO and members of the *LYR-IA* and *LYR-IIIA* groups have high affinity for LCO. However, no phylogenetic group in which members have conserved high affinity for short-chain CO has yet been identified.

The LYR and LYM proteins with well characterized biochemical properties were also studied for their roles in defence or symbioses. In non-legumes, the LCO receptor of the *LYR-IA* group plays a role in AM establishment^15,18^. However, knock out (KO) mutants were not completely deficient in their ability to be colonized by AMF, in contrast to mutants in the downstream CSSP, suggesting that additional receptors are involved. Since *LYR-IA* proteins are LCO receptors, it is thus expected that yet uncharacterized short-chain CO receptors play a complementary role to LCO receptors in CSSP activation.

AM has been lost in some plant species, such as the model plant *Arabidopsis thaliana*. This and other non-host plant species have lost some of the key genes required for this association such as genes of the CSSP^2^. Orthologs of genes missing in non-host plant species are thus likely to be involved in AM in host species.

Here we aimed to identify additional LysM-RLK involved in AM establishment and in particular the yet unknown high affinity short chain CO receptors.

## Results

### Petunia hybrida and Brachypodium distachyon LYR-IB LysM-RLKs have high affinity for LCO and CO

*LYR-IA* LysM-RLKs are LCO binding proteins only found in mycotrophic plant species and involved in AM establishment. The *LYR-IB* phylogenetic group is the closest to the *LYR-IA* group, and *LYR-IB* genes are also only present in mycotrophic plant species, generally represented by a single gene except in polyploid species (Suppl Figure 1). We hypothesized that members of this group are also involved in symbiotic signal perception and AM establishment.

In order to perform ligand binding assays with symbiotic signals, we used an *Agrobacterium tumefaciens*-mediated transient expression system to produce PhLYK9-YFP and BdLYR2-YFP in *Nicotiana benthamiana* leaves. We also expressed PhLYK15-YFP, a *P. hybrida* LysM-RLK belonging to another phylogenetic group (*LYR-IIB*), as a control. All proteins were found, by microscopy, in the cell periphery (Figure 1a) and immunodetected in the membrane fractions extracted from the corresponding *N*. *benthamiana* leaves (Figure 1b). Membrane fractions were used for ligand binding assays using either a radiolabelled LCO (LCO-V(C18:1,NMe,^35^S)) or a crosslinkable biotinylated CO5 (CO5-biot) and competitions with unlabelled LCO and CO. Binding of the radiolabelled LCO was detected in membrane fractions from leaves expressing PhLYK9-YFP and BdLYR2-YFP but not from leaves expressing PhLYK15-YFP or from untransformed leaves (Figure 1b). Affinity of PhLYK9 and BdLYR2 for LCO-V(C18:1,NMe,S) was measured by cold saturation experiments. Scatchard plot analysis revealed a single class of binding sites (Figure 1c) with dissociation constants (*K_d_*) of 2.8 nM± 0.8 nM (n=3) and 2.6 nM ± 1.1 (n=5), respectively. We then determined whether the PhLYK9 and BdLYR2 binding sites are specific for LCO through competition assays between the radiolabelled LCO and 1µM of unlabelled LCO, CO, chitosan (deacylated chitin) fragments or PGN fragments. CO4 and CO8 were efficient competitors of the radiolabelled LCO but not the chitosan or PGN fragments (Figure 1d). We further characterized the binding site of both proteins by competition assays between the radiolabelled LCO and a range of unlabelled CO4 or CO8 (Figure 1e) and determined inhibitory constants (*Ki*) between of 7.5 and 44.6 nM (Figure 1f).

**Figure 1.**
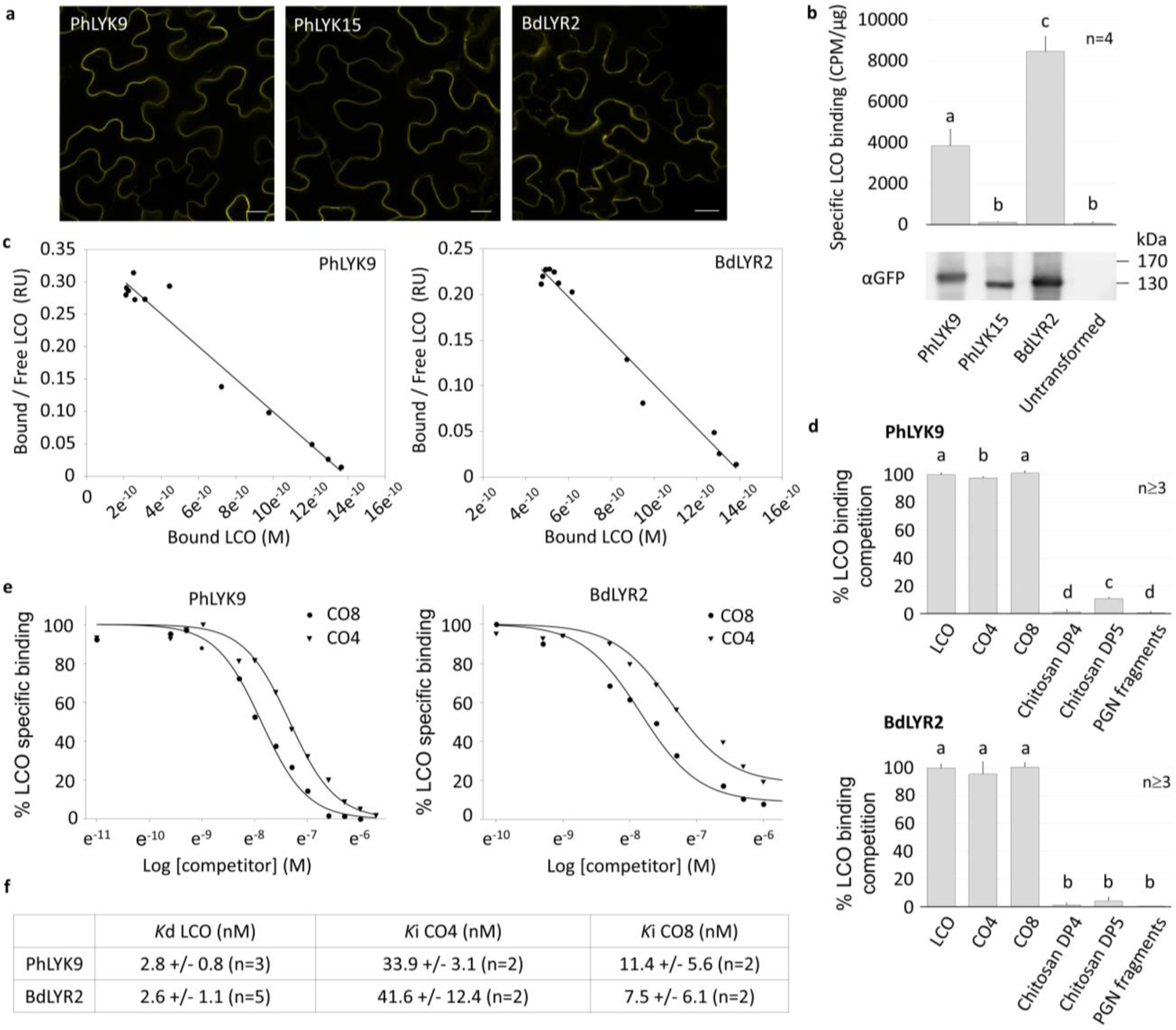
Equilibrium binding with a radiolabelled LCO shows high affinity LCO and CO binding to PhLYK9 and BdLYR2. **a**. Images of epidermal cells from *N*. *benthamiana* leaves expressing the indicated proteins tagged with YFP. Scale bars represent 20 µm. **b**. Specific binding of LCO-V(C18:1,NMe,^35^S) to PhLYK9 and BdLYR2. Bars represent the specific LCO-binding/µg membrane proteins (means and SD from two replicates on two independent batches of membrane fractions containing the indicated proteins or from untransformed leaves). Immunodetection with anti YFP antibodies in 10µg of the indicated membrane fractions. **c**. Affinity of PhLYK9 and BdLYR2 for LCO-V(C18:1,NMe,S). Scatchard plots of cold saturation experiments using membrane fractions containing the indicated proteins and a range of LCO-V(C18:1,NMe,S) concentrations as competitor. The plots are representative of experiments performed with two independent batches of membrane fractions. **d**. Specificity of PhLYK9 and BdLYR2 LCO-binding sites for LCO versus other GlcNAc-containing molecules. Bars represent the percentage competition of specific LCO-V(C18:1,NMe,^35^S) binding (means and SD between at least 3 technical replicates) in the presence of 2 μM of the indicated competitors. LCO is LCO-V(C18:1,NMe,S) **e**. Affinity of PhLYK9 and BdLYR2 for CO4 and CO8. Competitive inhibition of the radiolabelled LCO-V(C18:1,NMe,S) binding to membrane fraction containing the indicated proteins, using a range of concentrations of unlabelled CO4 or CO8 as competitors. **f**. Affinities of PhLYK9 and BdLYR2 LCO-binding sites for the indicated ligands, deduced from saturation and competitive inhibition experiments. Statistical differences (p-value < 0.05) were calculated using a pairwise T test in **a** and **d**.

Binding of CO5-biot (Figure 2a, Suppl Figure 2a), was observed on membrane fractions containing PhLYK9-YFP but not on those containing PhLYK15-YFP (Figure 2b). To confirm that CO5-biot was crosslinked to PhLYK9-YFP, we immunopurified PhLYK9-YFP and PhLYK15-YFP. CO5-biot was co-purified with PhLYK9 (Figure 2c). CO5-biot was also detected in membrane fraction containing BdLYR2 (Figure 2d). Affinity of PhLYK9-YFP and BdLYR2-YFP for CO5-biot was measured by saturation experiments. Saturation was achieved with 50 nM CO5-biot and dissociation constants (*K_d_*) were estimated around 25 nM (Figure 2e and g, Suppl Figure 3a-b). We performed competition assays between CO5-biot and a range of unlabelled LCO and also found inhibitory constants (*Ki*) estimated around 25 nM (Figure 2f and h).

**Figure 2.**
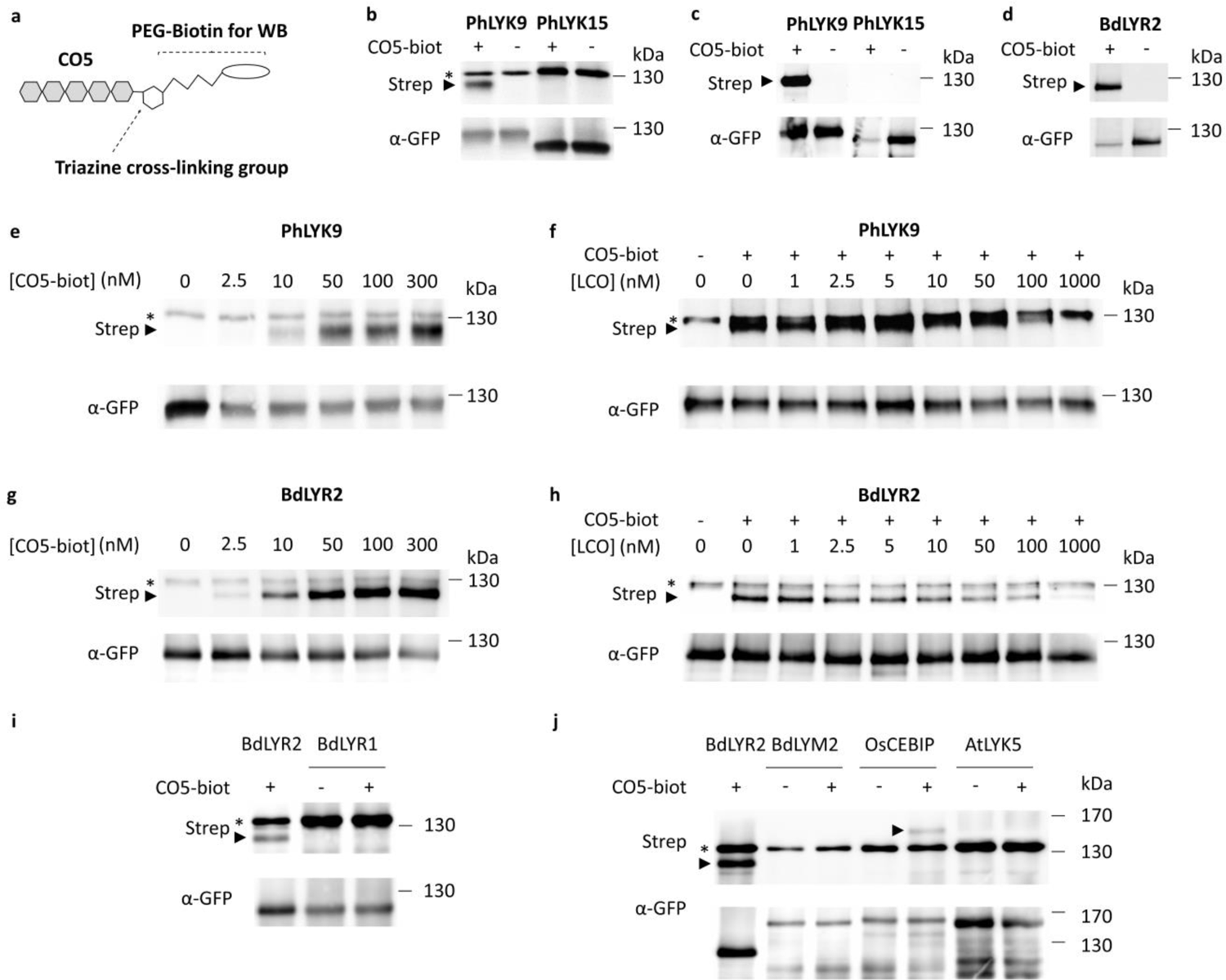
Crosslinking with a functionalized CO5 shows high affinity LCO and short-chain CO binding to PhLYK9 and BdLYR2. **a**. Schematic representation of the cross-linkable biotinylated CO5 (CO5-biot). The triazine group forms a covalent bound with adjacent proteins and the biotin-labeled protein can be detected by Western-blotting (WB). **b**. CO5-biot binding to 10 µg of *N. benthamiana* membrane proteins containing PhLYK9-YFP or PhLYK15-YFP. Membrane fractions were incubated with or without 10 nM CO5-biot and WB performed using sequentially anti-GFP antibodies and streptavidin on the same membrane. The arrowhead indicates the position of PhLYK9-YFP, whereas * indicates an *N. benthamiana* endogenously biotinylated protein. **c-d**. CO5-biot binding to immunopurified PhLYK9, PhLYK15 or BdLYR2. 250 µg of membrane proteins containing the indicated proteins were incubated with or without 1 µM nM CO5-biot, before protein solubilization and purification using anti-GFP beads. Note that the endogenously biotinylated protein detected in membrane fractions (**b**) was not detected anymore after protein purification. **e**,**g**. Affinity of PhLYK9 (**e**) and BdLYR2 (**g**) for short-chain CO. Saturation experiments on 10 µg of membrane proteins using a range of concentrations of CO5-biot. **f**,**h**. Specificity of the PhLYK9 (**f**) and BdLYR2 (**h**) CO-binding sites for CO versus LCO. Ten µg of membrane proteins were incubated with or without 10 nM CO5-biot and a range of unlabelled LCO-V(C18:1,NMe,S). **i-j**. CO5-biot binding to 5 µg and 25 µg of membrane proteins containing indicated proteins. Membrane fractions were incubated with or without 10 nM CO5-biot.

The two *LYR-IB* proteins, PhLYK9 and BdLYR2 each have at least one binding site with high affinity for both LCO and CO, with no selectivity between LCO and CO. In contrast, *LYR-IA* proteins have a high affinity LCO binding site that is selective for LCO versus CO^12,15^. To exclude that *LYR-IA* proteins have an additional high affinity short-chain CO binding site, we measured CO5-biot binding to BdLYR1. No binding was detected to this protein, in comparison to BdLYR2 (Figure 2i). LysM-RLK/Ps belonging to other phylogenetic groups are known to have high affinity for long-chain CO^8,10^. We measured CO5-biot binding to the rice OsCEBIP (*LYM-II* group), its ortholog in *B. distachyon*, BdLYM2, and the *A. thaliana* AtLYK5 (*LYR-IIIC* group). Only a faint signal was detected for OsCEBIP and no signal was detected for BdLYM2 and AtLYK5, in comparison to BdLYR2 (Figure 2j).

As an alternative method to produce the proteins, we expressed ECRs of BdLYR2 and BdLYR1 in insect cells and used MicroScale Thermophoresis (MST) to perform ligand binding assays. Pure BdLYR1 and BdLYR2 proteins were obtained by FPLC (Figure 3a). We labelled the proteins with a red fluorophore and performed MST using a range of concentrations of unlabelled LCO or CO (Figure 3b). For BdLYR2, similar ranges of affinity were determined for LCO, CO4, CO5 and CO8 (Figure 3c-d, Suppl Figure 4). For BdLYR1, affinity for CO4 was much lower than for LCO (Figure 3e-f, Suppl Figure 4). We also performed MST with unlabelled proteins and a fluorescent CO5 (Figure 3g) and observed higher affinity of BdLYR2 than BdLYR1 for the CO5-BODIPY (Figure 3h-i). Despite the selectivity of BdLYR1 and BdLYR2 for LCO and CO being the same as those obtained with full-length proteins expressed in *N. benthamiana* leaves, the affinities found with ECR expressed in insect cells were much lower (µM range versus nM range).

**Figure 3.**
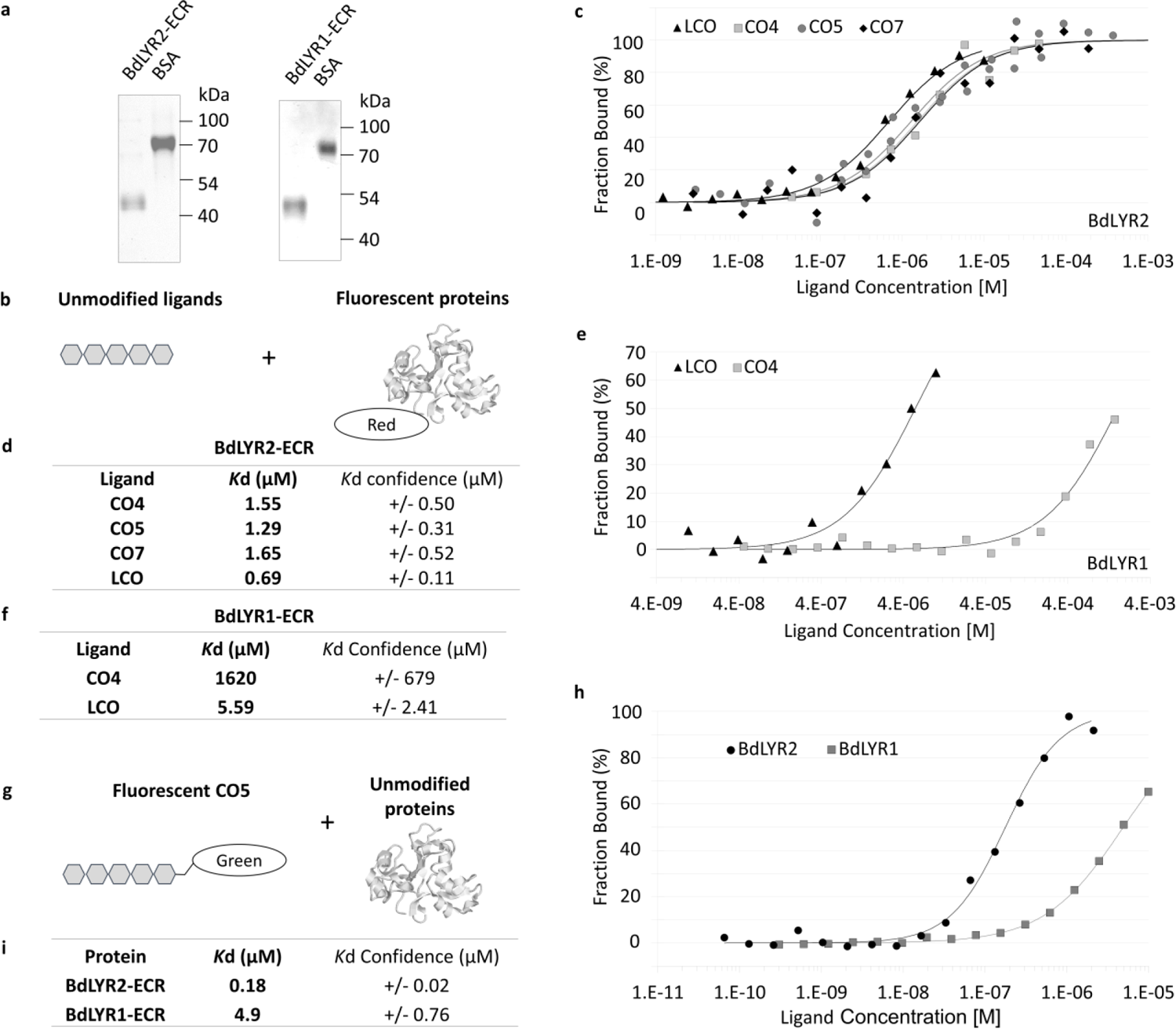
Microscale thermophoresis confirms the absence of selectivity of BdLYR2 between LCO and CO. **a**. SDS-PAGE and coomassie blue staining of purified BdLYR2-ECR or BdLYR1-ECR and 400 ng or 200 ng of BSA respectively. **b**. Schematic representation molecules used for MST: Unlabelled ligands and ECR labelled with a red fluorophore. **c**. Curves of LCO and CO binding to BdLYR2-ECR. 20nM of BdLYR2-ECR were incubated with a range of LCO-V(C18:1,NMe,S), CO4, CO5 or CO7. The plots show the fraction bound (mean ΔF_norm_ values divided by the curve amplitude), from 2 independent experiments. Plots showing the binding amplitudes (ΔF_norm_) are in Suppl Figure 4. **d**. Affinity of BdLYR2-ECR for LCO and CO, deduced from the binding curves. **e**. Curves of LCO and CO binding to BdLYR1-ECR. 20nM of BdLYR1-ECR were incubated with a range of LCO-V(C18:1,NMe,S) or CO4. Plots with binding amplitudes are shown in Suppl Figure 4. **f**. Affinity of BdLYR1-ECR for LCO and CO, deduced from the binding curves. Note that since the curves with BdLYR1 are not saturated, the *K_d_* might be overestimated. **g**. Schematic representation molecules used for MST: fluorescent CO5-BODIPY and unlabelled ECR. **h**. Curves of BdLYR1-ECR and BdLYR2-ECR binding to CO5. 100nM of CO5-BODIPY were incubated with a range of BdLYR1-ECR or BdLYR2-ECR. Because CO5-BODIPY fluorescence changed upon binding to ECRs, fluorescence intensity rather than thermophoresis was analysed. The plots show the fraction bound (fluorescence intensity values divided by the curve amplitude), **f**. Affinity of BdLYR1-ECR and BdLYR2-ECR for CO5, deduced from the binding curves. Note that since the curves for BdLYR1-ECR in h are not saturated, the *K_d_* might be overestimated.

### LYR-IB proteins have conserved biochemical properties in other plants species

We found high affinity of the *M. truncatula* MtLYR8 for LCO (*K_d_* of 12.3 nM ± 0.3 nM, n=2) and short-chain CO (*K_d_* of about 25 nM) and absence of specificity for LCO versus CO (Suppl Figure 5a-d). In the hexaploid wheat, the three *LYR-IB* proteins *TaLYR2A*, *TaLYR2B* and *TaLYR2D* showed LCO binding and absence of selectivity between LCO and CO (Suppl Figure 5e-f). We then characterized in more detail TaLYR2D and found high affinity for LCO (*K_d_* of 1.5 nM ± 0.4nM, n=2) and short-chain CO (*K_d_* of about 25 nM) (Suppl Figure 5g-i).

Altogether, these results show that high affinity for short-chain CO is only for members of the *LYR-IB* group and not for members of the *LYR-IA*, *LYR-IIIC* and *LYM-II* groups, which respectively have high affinity for LCO (*LYR-IA*)^12, 14–16^ and long-chain CO (*LYR-IIIC* and *LYM-II*)^8–11^.

### LYR-IB LysM-RLKs are involved in AM establishment in Petunia hybrida and Brachypodium distachyon

We identified a KO line in the *P. hybrida LYR-IB* coding sequence (*Phlyk9-1*, Figure 4a). *Phlyk9-1* homozygous plants displayed a reduced number of AMF colonization sites at early stages of colonization (Figure 4b) and a reduced root length colonization by AMF at later stages of colonization (Figure 4c-d), compared to segregating plants with the wild type (WT) *PhLYK9* allele. We also produced a CRISPR/Cas9 KO line in the *LYR-IB* gene of a closely related species, tomato (*Sllyk9-1*, Suppl Figure 6a-b). A reduced colonization by AMF was also observed in this line (Suppl Figure 6c). A KO line in the *P. hybrida LYR-IA* gene (*Phlyk10-1*) also showed a decrease in AMF colonization in Girardin et al. (2019)^15^. We thus crossed *Phlyk9-1* and *Phlyk10-1* to test potential redundancy between the genes. We observed a reduced AMF colonization in the double mutant compared to the single *Phlyk9-1* mutant (Figure 4e).

**Figure 4.**
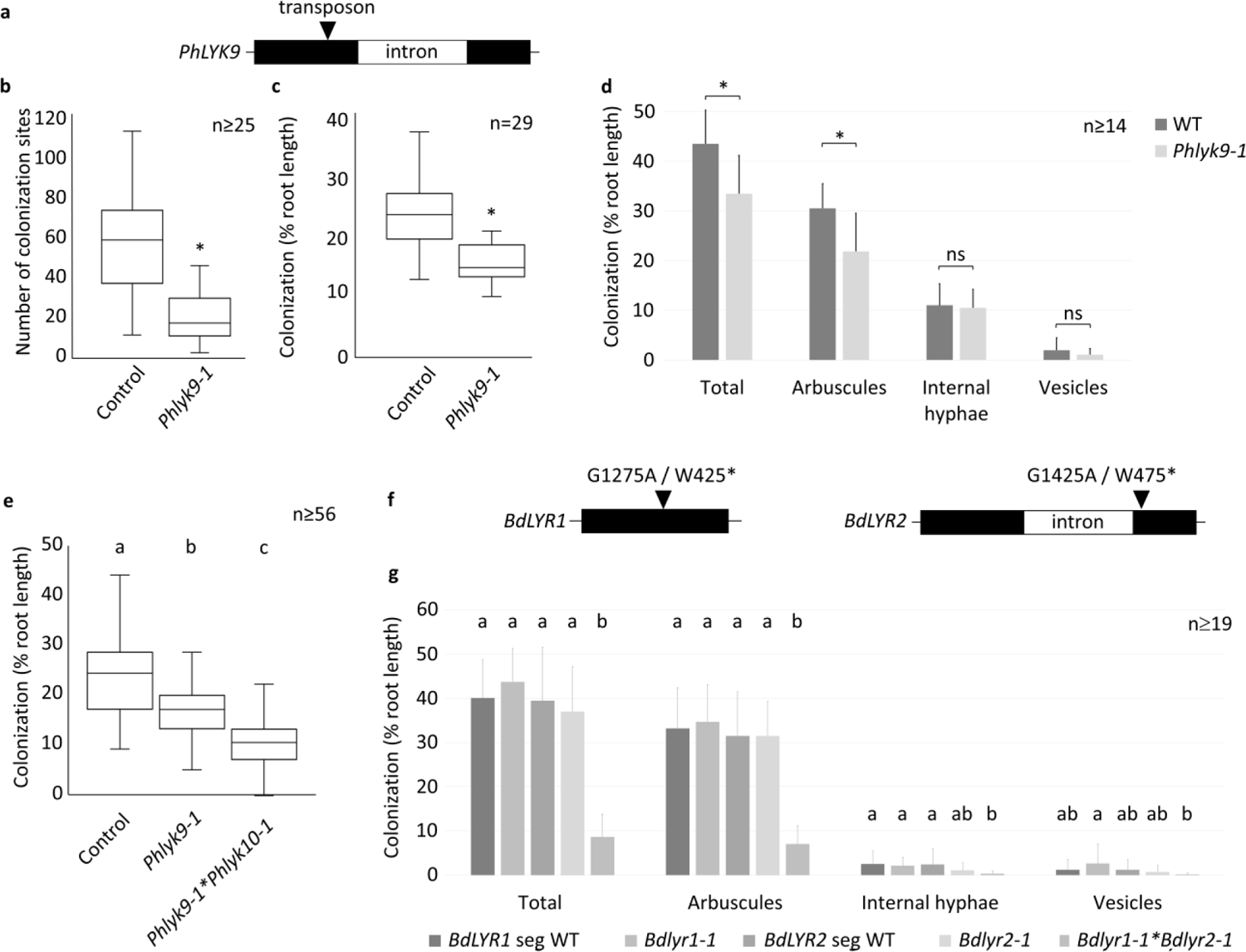
*PhLYK9* and *BdLYR2* are involved in AM establishment. **a**. Position of the dTPh1 transposon insertion in *Phlyk9-1*. **b**. Number of AMF colonization sites per root system at 3 wpi in segregating plants bearing homozygous *Phlyk9-1* or WT *PhLYK9* (control) alleles. Boxplots represent the distribution between root systems from 2 independent experiments. **c**. Root-length colonization at 4 wpi in segregating plants bearing homozygous *Phlyk9-1* or WT *PhLYK9* alleles. Boxplots represent the distribution between root systems from 3 independent experiments. **d**. Detailed analysis of AMF structures at 6 wpi in segregating plants bearing homozygous *Phlyk9-1* or WT *PhLYK9* alleles. Means and SD between root systems from 2 independent experiments are shown. **e**. Root-length colonization at 4 wpi in segregating plants bearing homozygous *Phlyk9-1* and *Phlyk10-1, Phlyk9-1* and WT *PhLYK10* (*Phlyk9-1*) or WT *PhLYK9* and *PhLYK10* (control) alleles after the cross of *Phlyk9-1* and *Phlyk10-1* lines. Boxplots represent the distribution between root systems from 3 independent experiments. **f**. Positions of the nonsense mutations in *Bdlyr1-1* and *Bdlyr2-1*. **g**. Detailed analysis of AMF structures at 4 wpi in segregating plants bearing homozygous *Bdlyr-1* or WT *BdLYR1* (Seg WT) and segregating plants bearing homozygous *Bdlyr2-1* and WT *BdLYR1* (*Bdlyr2-1*), *Bdlyr2-1* and *Bdlyr1-1*, or WT *BdLYR2 and* WT *BdLYR1* (Seg WT) after the cross of *Bdlyr2-1* and *Bdlyr1-1* lines. Means and SD between root systems from 2 independent experiments are shown. Statistical differences (p-value < 0.05) were calculated using a Wilcoxon test in **b** and **c** and a Kruskal Wallis test in **d**, **e** and **g**.

We also identified KO lines in the *B. distachyon LYR-IA* and *LYR-IB* coding sequences (*Bdlyr1-1* and *Bdlyr2-1,* respectively, Figure 4f). In contrast to *Phlyk10-1* and *Phlyk9-1*, neither *Bdlyr1-1* nor *Bdlyr2-1* homozygous plants showed differences in root length colonization by AMF (Figure 4g) compared to their segregating WT plants. However, the *Bdlyr1-1/Bdlyr2-1* double mutant showed a strong decrease in colonization by AMF (Figure 4g), as found for *Phlyk10-1/Phlyk9-1*.

### The M. truncatula LYR-IB LysM-RLK controls AM and LCO and short-chain CO responses

We used the legume *M. truncatula* to analyse the role of the *LYR-IB* gene in both AM and nodulation and in activation of root-epidermal Ca responses to LCO and short chain CO, which have been well characterized in this species^4–5^. Two KO lines (*Mtlyr8-1* and *Mtlyr8-2)* were produced by CRISPR/Cas9 gene editing and *A. tumefaciens* transformation (Figure 5a, Suppl Figure 7a-b). A reduced colonization by AMF was found in the two lines (Figure 5b; Suppl Figure 7c). In contrast, both mutant lines nodulated at least as well as WT with both the WT *M. truncatula* Rhizobial partner *Sinorhizobium meliloti* and the *nodFE* strain (Suppl Figure 7d-e), the latter of which produces LCO more similar to the ligand used for binding. We then used *Agrobacterium rhizogenes* transformation to introduce the Ca concentration GECO reporter and measured calcium spiking responses in root atrichoblasts following CO4 or LCO treatments. The LCO used (LCO-V(C18:1,NMe,S) as for the ligand binding assays, has a structure different from those produced by *S. meliloti* (LCO-IV(C16:2,Ac,S)), and thus is expected to produce responses more related to AM than nodulation. Both LCO and short-chain CO induced calcium spiking were strongly reduced in the transgenic roots of the two KO lines compared to the controls (Figure 5c, Suppl Figure 7f), thus clearly showing a role of the *LYR-IB* receptor in symbiotic LCO and short-chain CO responses.

**Figure 5.**
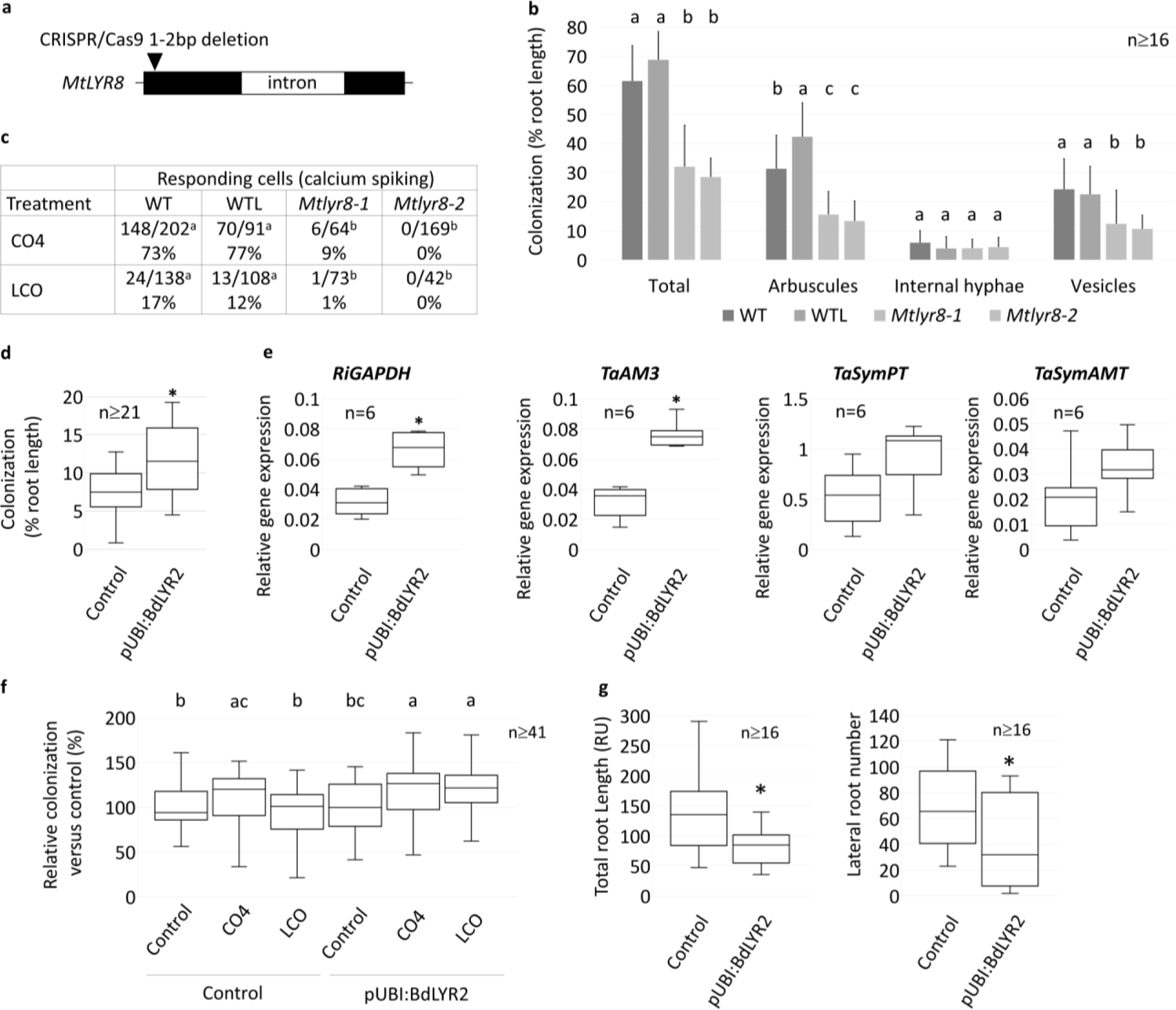
*MtLYR8* controls LCO and short-chain CO responses and AMF colonization in *M. truncatula* and strong expression of BdLYR2 in wheat affects AMF colonization and root architecture. **a**. Position of the frameshift deletions in *Mtlyr8-1* and *Mtlyr8-2*. Details are shown in Suppl Figure 7b. **b**. Detailed analysis of AMF structures at 4 wpi in *Mtlyr8-1, Mtlyr8-2,* 2HA (WT) or a 2HA line transformed with an empty vector (WTL). Means and SD between root systems from 2 independent experiments. **c**. Number and proportion of root atrichoblast cell nuclei showing calcium spiking to all fluorescent nuclei analysed following treatment with 10^-7^ M CO4 or LCO-V (C18:1,NMe,S). Data are the sum from at least 3 independent plants. Only nuclei with two or more spikes in 20 min were considered as responding cells. **d**. Root-length colonization at 32 dpi in wheat plants expressing BdLYR2 under the control of a strong promoter (pUbi:BdLYR2) or containing the empty vector (Control). Boxplots represent the distribution between root systems of 2 independent transgenic lines for each construct. **e**. Relative expression of the *R. irregularis* gene *RiGAPDH* (reflecting the amount of fungus in the roots), the early AM stage marker genes TaAM3 and the ammonium and phosphate transporters specifically expressed in arbuscule-containing cells (SymAMT2 and symPT respectively) in pUbi:BdLYR2 and control plants at 6 wpi. Boxplots represent the distribution between 3 pools of roots from two independent transgenic lines for each construct. **f**. Effect of LCO and CO on root-length colonization in wheat plants expressing BdLYR2 under the control of a strong promoter (pUbi:BdLYR2) or containing the empty vector (Control). Plants were treated with with 10^-7^ M CO4 or LCO-V(C18:1,NMe,S). Boxplots represent the distribution between root systems of 2 independent transgenic lines for each construct and of 2 independent experiments (harvested at 5 or 6 wpi). Relative colonization levels to mean colonization in non-treated control plants in each experiment. **g**. Effect of pUbi:BdLYR2 on root architecture. Boxplots represent the distribution between root systems of 2 independent transgenic lines for each construct. Statistical differences (p-value < 0.05) were calculated using a Van Waerden test in **b**, a Chi-square test in **c**, T tests in **d** and **e** and a Kruskal Wallis test in **f** and **g**.

### Strong expression of a LYR-IB gene increases AMF colonization in wheat

Since we showed that *LYR-IB* proteins are involved in symbiotic signal perception and AM establishment, we tested whether expression under the control of a strong constitutive promoter could increase colonization by AMF. We chose *Triticum aestivum*, as a major crop, for these experiments. We produced transgenic wheat lines expressing *BdLYR2* under the control of a strong promoter. We found an increase in root colonization by AMF of transgenic plants compared to controls when the colonization level was low (Figure 5d), but this effect was not observed at higher levels of colonization (Suppl Figure 8a). Using molecular markers to quantify AMF colonization, we found a significant increase in the transcript level of an AMF gene and a wheat AM marker gene (Suppl Figure 8b), in transgenic plants compared to controls (Figure 5e). However, although their expression was increased, the transcript levels of two transporters specifically expressed in arbuscule-containing cells were not significantly different in transgenic plants compared to controls (Figure 5e). To further characterize the transgenic plants, we quantified the stimulation of AMF colonization by LCO and CO4 treatments. Treatment with 10^-7^ M CO4, but not 10^-7^ M LCO, increased AMF colonization in the control plants. In contrast to the control plants, treatment with 10^-7^ M LCO was able to increase AMF colonization in the transgenic plants while no further increase was seen compared to control plants with CO4 (Figure 5f).

Finally, as LysM-RLKs are known to control root development in response to chitinic signals^19^, we measured root architecture in the transgenic wheat plants grown in axenic conditions, and found that both total root length and number of lateral roots were reduced, in comparison to the control plants (Figure 5g).

### Atypical evolution of the LYR-IB gene in rice

*LYR-IB* genes were found in all angiosperm species we looked at (Suppl Figure 1)^7^ and had conserved biochemical properties and roles in AM establishment in the species we have studied. However, we found the full coding sequence of the *LYR-IB* gene in indica rice genomes but not in the two japonica reference genomes, Nipponbare and Kitaake. In these two genomes, there is a one bp deletion in the first exon that creates a frameshift (Figure 6a), leading to a predicted protein truncated just before the TM. Such a protein would not anchor to the PM, and thus is probably mislocalised and non-functional in signal transduction. To determine whether this deletion was specific to these two japonica cultivars, we sequenced an amplicon covering the deletion in 15 japonica and 26 indica cultivars (Suppl Figure 9). The deletion was absent in all indica cultivars. In contrast the deletion was present in most japonica cultivars. We verified that the full-length *LYR-IB* protein from the indica cultivar Xihui18 (*OsLYK11*) has similar biochemical properties to the other *LYR-IB* proteins. We found that OsLYK11-YFP, produced in *N. benthamiana* leaves, has high affinity for both LCO (*K_d_* of 4.7 nM ± 1.7 nM, n=2) and short-chain CO (*K_d_* of about 25 nM) (Figure 6b-d). CO5-biot binding was detected for OsLYK11 but not for the *LYR-IA* OsNFR5/OsMYR1 (Figure 6e). A CRISPR/Cas9 Xihui18 KO line for *OsLYK11* (*Oslyk11-1*, Figure 6f, Suppl Figure 10a-b) showed no difference in root colonization by AMF compared to WT plants (Figure 6g). Since japonica cultivars are natural *OsLYK11* mutants, we introduced the indica Xihui18 *OsLYK11* gene in Kitaake and Nipponbare and quantified AMF colonization. Again, no differences were found between the control plants and those containing the indica *OsLYK11* allele (Figure 6h, Suppl Figure 10c).

**Figure 6.**
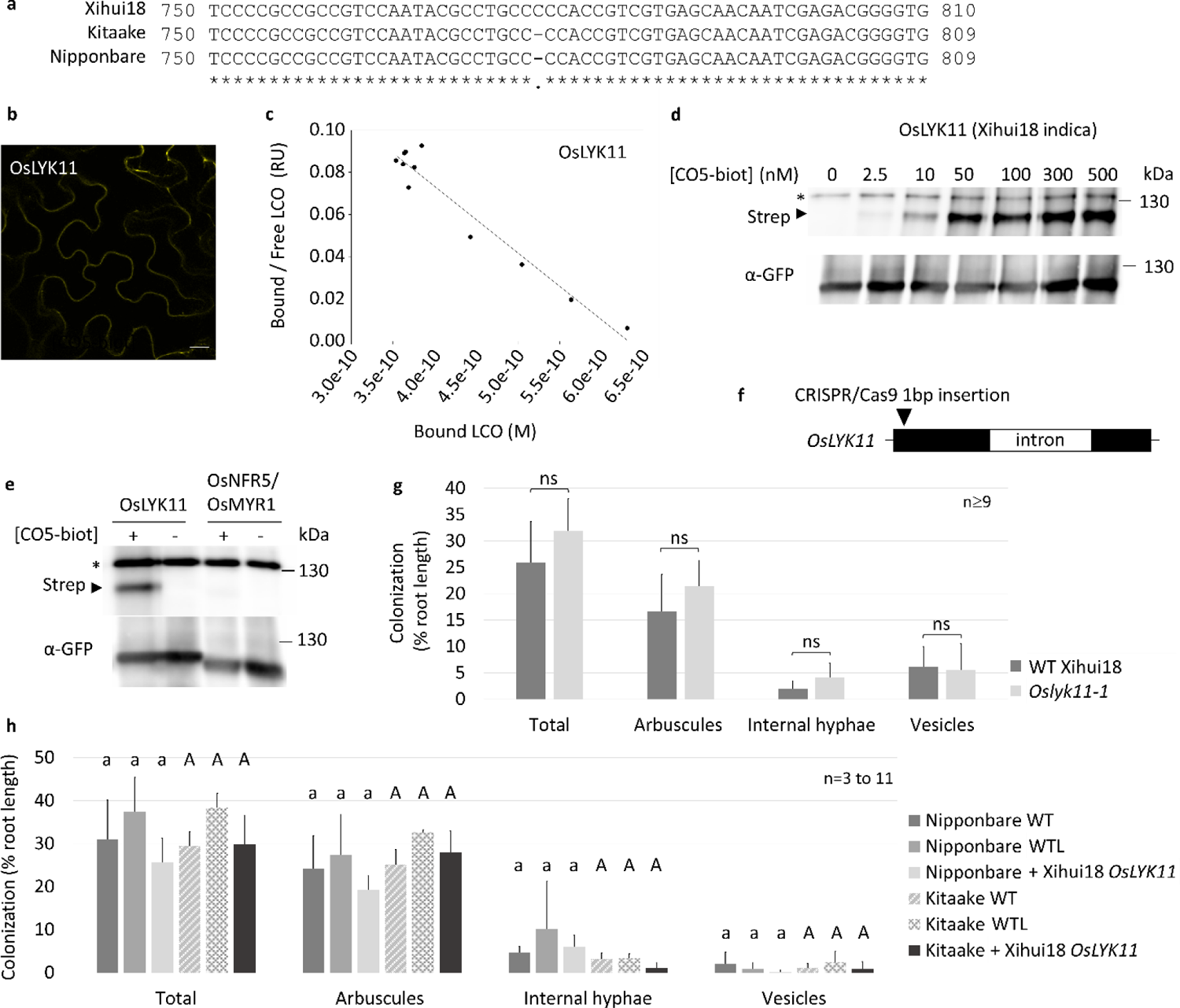
*OsLYK11* is conserved in indica but not in japonica rice. **a**. Partial sequence alignment of *OsLYK11* from the indica rice cultivar Xihui18 and the japonica rice cultivars Nipponbare and Kitaake, showing the natural 1 bp deletion in the japonica cultivars. **b**. Image of epidermal cells from a *N*. *benthamiana* leaf expressing Xihui18 OsLYK11-YFP (OsLYK11). Scale bar represents 20 µm. **c**. Affinity of OsLYK11 for LCO-V(C18:1,NMe,S). Scatchard plot of a cold saturation experiment using membrane fractions containing OsLYK11 and a range of LCO-V(C18:1,NMe,S) concentrations as competitor. The plot is representative of experiments performed with two independent batches of membrane fractions. **d**. Affinity of OsLYK11 for short-chain CO. Saturation experiments on 10 µg of membrane proteins containing OsLYK11 using a range of concentrations of CO5-biot. WB performed using sequentially anti-GFP antibodies and streptavidin on the same membrane. The arrowhead indicates the position of OsLYK11-YFP, whereas * indicates an *N. benthamiana* endogenously biotinylated protein. **e**. CO5-biot binding to 10 µg of *N. benthamiana* membrane proteins containing OsLYK11 or OSNFR5/OsMYR1. **f**. Position of the frameshift insertion in Xihui18 *Oslyk11-1*. Details are shown in Suppl Figure 10b. **g**. Detailed analysis of AMF structures at 5 wpi in *Oslyk11-1* or WT Xihui18 plants. Means and SD between root systems from 2 independent experiments are shown. **h**. Detailed analysis of AMF structures at 5 wpi in Nipponbare or Kitaake plants WT, containing the empty vector (WTL) or containing Xihui18 *OsLYK11*. Means and SD between roots systems of 2 independent transgenic lines and 1 WT line from 1 experiment. A second replicate is shown in Suppl Figure 10c. Statistical differences (p-value < 0.05) were calculated using a Van Waerden test in **g** and **h**.

## Discussion

### LYR-IB LysM-RLKs are LCO and short-chain CO receptors

Short-chain CO are important signalling molecules produced by AMF that elicit symbiotic responses at nM concentrations. By characterising members of the *LYR-IB* phylogenetic group in *P. hybrida*, *M. truncatula*, *B. distachyon*, rice and wheat, we have identified a clade of LysM-RLKs which show high affinity for short chain CO, in addition to LCO and longer CO. Among the LysM-RLKs characterized to date, these *LYR-IB* proteins are the only ones described to have high affinity for short-chain CO. In comparison, several LysM-RLPs of the *LYM-II* group and the LysM-RLK of the *A. thaliana LYR-IIIC* group, AtLYK5, have high affinity for long-chain, but not to short-chain CO (this work and ^9,10,12,20^) and although the rice *LYR-IA* OsNFR5/OsMYR1 was described to have high affinity for short-chain CO^18^, we found lower affinity for short-chain CO in this protein than in *LYR-IB* proteins.

We compared several expression systems and ligand binding assays. We found similar affinities (nM range) with the two ligand binding assays (radiolabelled LCO or crosslinkable CO5) when we used full-length proteins produced in *N. benthamiana* leaves, but much lower affinities (µM range) when we used the ECRs expressed in insect cells combined with MST. Low affinity (µM range) was also found for OsCEBIP ECR produced in insect cells^11^ while binding assays on the full-length proteins showed high affinity (nM range)^20^, coherent with characterized endogenous high affinity (nM range) binding sites^21^. The discrepancy between the data we obtained with ECRs and full-length proteins might result from the ability of full-length proteins to form homo-oligomers (or hetero-oligomers with *N. benthamiana* endogenous proteins) leading to increased affinity for the ligand. Despite the discrepancy in affinity, selectivity for the ligand was conserved between the two expression systems tested, and confirmed the ability of *LYR-IB* proteins to bind both short-chain CO and LCO. It however highlights the caution required to compare data from the literature obtained with various expression systems/ligand binding assays.

The biochemical properties we found for full-length *LYR-IB* proteins are coherent with an endogenous short-chain CO binding site described in a tomato cell culture and derived membrane fraction using a radiolabelled CO5^22^. In the membrane fractions, this site had a *K_d_* of 22.8 nM for their modified CO5 and did not discriminate CO4, CO5 and a LCO-V(C18:1,Ac). These characteristics were consistent with the concentration of CO4 and CO5 inducing biological activities^23^ suggesting that it is the major short chain CO binding site in the tomato cell culture. The function of *LYR-IB* proteins as major short-chain CO and LCO receptors is further shown by the large decrease in short-chain CO and LCO signalling (decrease in calcium spiking) we measured in roots of *M. truncatula Mtlyr8* and that has also been found in the barley *LYR-IB* mutant, *Hvrlk2*^5^.

The absence of selectivity between CO4/5, CO8 and LCO of *LYR-IB* raises the question of which structure is recognized by the proteins. It is possible that the binding pocket accommodates 4 GlcNAc that are sufficient for high affinity binding, and that additional GlcNAc (in longer chain CO) or an acyl chain (in LCO) do not interfere with the binding affinity.

Various chitinic molecules including short-chain CO and LCO can induce lateral root formation in various plant species including in dicots and monocots^19,25,26^. This phenomenon was shown to rely on *LYR-IA, LYM-II* and *LYK-I* LysM-RLKs and at least a part of the CSSP^19,25^. The impact on wheat root architecture of ectopic *BdLYR2* expression under a strong promoter, further supports the role of LysM-RLKs in root developmental processes and extends it to the *LYR-IB* LysM-RLKs. Strong expression of LysM-RLKs might induce signalling in the absence of a ligand, as observed with spontaneous nodulation in legumes^15,27^ or activation of cell death in *N. benthamiana*^28^.

### Multiple symbiotic signal receptors are involved in AM establishment

We previously showed that Solanaceae (petunia and tomato) mutants in *LYR-IA* LCO receptors have a decreased colonization by AMF. Here, we show that Solanaceae (petunia and tomato) and Fabaceae (*M. truncatula*) mutants in *LYR-IB* CO / LCO receptors have also a decrease in colonization by AMF, and that in petunia the double mutant has a more severe AM phenotype than the single *LYR-IB* mutant. In *B. distachyon* (this study) and in barley^5^, single *LYR-IA* and *LYR-IB* mutants do not have AM phenotypes, and only double mutants have severe phenotypes, as in the petunia double mutant, suggesting different levels of *LYR-IA* and *LYR-IB* redundancy in monocots compared to dicots. Interestingly both *LYR-IA* and *LYR-IB* gene expression were under the control of nitrogen and phosphate deficiency in barley, but not in *M. truncatula*^5^. However, in *M. truncatula*, *LYR-IB* appears to have retained an ancestral role in AM, and has not evolved a dual role in nodulation with rhizobial bacteria as *LYR-IA*, which is consistent with poor *LYR-IB* protein binding to *S. meliloti* LCO^12^ compared to the LCO used here.

Because of the absence of a functional *LYR-IB* gene in the rice cultivar Nipponbare, the AM phenotype found in *LYR-IA* mutants^18^ is thus equivalent to the *LYR-IA*/*LYR-IB* double mutants in other Poaceae. However, we found by knocking out the *LYR-IB* gene in the indica Xihui18 cultivar and by complementing the natural *LYR-IB* gene KO in the japonica Nipponbare and Kitaake cultivars, that a single *LYR-IB* mutant in rice has no AM phenotype, as in other Poaceae. The deletion in the *LYR-IB* gene in most japonica rice is reminiscent of the situation with *OsCERK1* (a LysM-RLK belonging the *LYK-I* group), for which haplotypes from japonica cultivars show lower efficacy for AM establishment than those from indica cultivars^29^ suggesting different selection pressures on AM related genes in domestication/breeding of indica and japonica rice.

In conclusion, this work has identified that the *LYR-IB* LysM-RLKs directly bind short-chain CO and LCO signals and play an important role in signalling and AM establishment.

## Materials and Methods

### Mutant lines

The *Phlyk9-1* mutant allele (line LY3182, dTph1 insertion 718bp from the start codon, W118 genotype) was identified by BLAST search in a Petunia dTPh1 transposon flanking sequence database^30^ with the full PhLYK19 coding sequence. *Phlyk10-1* is described in^15^. Progeny genotyping was performed by PCR using the primers listed in Suppl data.

Induced mutations in *Bdlyr1-1* (line 5525) and *Bdlyr2-1* (line 5160) were isolated from an sodium azide-mutant population of *Brachypodium distachyon Bd21-3* genotype as described^31^ on the EPITRANS platform^32^. The mutation detection system used in TILLING is based on NGS sequencing of libraries of PCR fragments, labeled with unique sequences and amplified from pooled samples in several dimensions^33^. The mutant lines are then identified using the SENTINEL software^34^.

Progeny genotyping was performed by dCAPS (endogenous *HaeIII* restriction site (*BdLYR1*) or site created with the primer (*BdLYR2*) lost in the mutant alleles) and confirmed by PCR amplicon sequencing with the primers listed in Suppl data.

*Sllyk9-1* was produced by *A. tumefaciens* transformation of the Ailsa Craig cultivar by CRISPR/Cas9 gene editing using the vector described in^35^ and a protospacer described in Suppl Figure 6. T2 plants were phenotyped.

*Mtlyr8-1* and *Mtlyr8-2* are independent lines produced by CRISPR/Cas9 gene editing via *A. tumefaciens* transformation of the 2HA genotype as described in^36^, using a protospacer described in Suppl Figure 7. A transformed non-edited line that segregated out the T-DNA was selected as control (WTL). T1 plants were phenotyped in Figure 5 and T2 plants in Suppl Figure 7.

*OsLyk11-1* was produced by CRISPR/Cas9 gene editing via *A. tumefaciens* transformation of the Xihui18 cultivar with the vector described in^35^ and a protospacer described in Suppl Figure 10. T1 plants were phenotyped.

### Transgenic lines

The rice Nipponbare and Kitaake cultivars were transformed by *A. tumefaciens* containing the plasmid PTCK303 in which a synthesized DNA fragment with the Xihui18 *pOsLYK11:OsLYK11* sequence modified to remove the restriction sites (sequence in Suppl data) was cloned between the *HindIII* and *BstEII* sites and a codon usage optimized *NLS-gGECO* (sequence in Suppl data) was cloned under the control of *pZmUBI* between the *KpnI* and *SacI* sites. Lines transformed with the same vector containing only the *pZmUBI:NLS-gGECO* were used as control. T1 plants were phenotyped.

The wheat Fielder cultivar was transformed by *A. tumefaciens* carrying the plasmid pBIOS12366 containing the following cassette: *pZmUBI:BdLYR2-cmyc*, *pZmUBI:NLS-gGECO*, and *pBradi1g20260:GUS* (sequences in Suppl data). Lines transformed with the plasmid pBIOS11613 containing only the *pZmUBI:NLS-gGECO* and *pBradi1g20260:GUS* (introduced as reporters for LCO and CO perception) were used as control. Note that the reporters were not detected either in absence or presence of LCO or CO. T0 lines with a single T-DNA insertion were selected by qPCR. T2 plants were phenotyped.

The NLS-gGECO and DsRed construct^37^ were introduced by *A. rhizogenes* in roots of the *M. truncatula* 2HA lines described above.

### In planta expression

For *AtLYK5, PhLYK9, BdLY2, TaLYR2A, TaLYR2B, TaLYR2D*, we cloned the coding sequences optimized with *N. benthamiana* codon usage (sequences in Suppl data) by gene synthesis, in translational fusion with the sequence coding *YFP* under the control of *Pro35S* in pCambia 2200 vector modified for Golden gate cloning^38^.

We also first cloned *PhLYK15, MtLYR8, OsLYK11* as described above (gene synthesis except for *PhLYK15* which was amplified by PCR from genomic DNA) but because their expression levels were too low or the proteins were mislocalized, the sequences encoding their ECR were amplified by PCR (primers listed in Suppl data) and cloned in translational fusion with the sequences coding with the MtNFP TM and ICR, and YFP under the control of *Pro35S* as described in^38^. We previously demonstrated that such chimeras have similar biochemical properties as the full-length proteins^12,15,38^.

In order to have a tagged OsCEBIP, the sequence encoding its ECR (optimized sequences in Suppl data) was also cloned in translational fusion with sequences coding MtNFP TM and ICR, and YFP as described above. Cloning of *OsNFR5/OsMYR1* and *BdLYR1* is described in^12^.

Leaves of *N. benthamiana* were infiltrated with the *A. tumefaciens* LBA4404 virGN54D strains. Leaves were harvested 3 days after infiltration.

### Insect cell expression

Synthetic DNA fragments containing coding sequence of the hemolin signal peptide, BdLYR2-ECR or BdLYR1-ECR without signal peptides, 3HA tag, PreScission protease cleavage site and 6HIS tag in translational fusion (sequences in Suppl data) were cloned in the pGTPb302 plasmid.

BdLYR1 was expressed in High Five cells at 27°C for 72 h (250 ml of cells (1.106 c/ml) infected with 0.536 µl of baculovirus (1.4 10^9^ qpfu/ml)). BdLYR2 was expressed in Sf9 cells at 27°C for 48 h (250 ml of cells (1.5 10^6^ c/ml) infected with 0.625 µl of baculovirus (3.94 10^9^ qpfu/ml).

The cells were centrifuged at 2000 *g* for 20 min at 4°C. The supernatant was diluted to half with 40 mM BisTris pH 6.8 and loaded onto HiPrep™ Q FF 16/10 anion exchange chromatography column (GE healthcare), washed with 20 mM BisTris pH 6.8 - 100 mM NaCl.

The bound proteins were eluted with 20 mM BisTris pH 6.8, 500 mM NaCl and further purified on histidine-tagged protein columns. HisTrap FF Crude (Cytiva) was used for BdLYR1 that was eluted in 20 mM Na phosphate pH 8, 500 mM NaCl, 40 mM imidazole and HiTrap Excel (GE healthcare) was used for BdLYR2 that was eluted in 50 mM Na phosphate pH8, 300 mM NaCl, 40 mM imidazole.

The purified proteins were concentrated with a Vivaspin 20 PES membrane unit with a 10kDa cut off, and buffer was exchanged with 50 mM Tris pH 7.4, 150 mM NaCl, using a *spin column A buffer exchange* (Monolith NT Protein Labelling Kit RED-NHS) before storage at −80°C.

### AM phenotyping

Petunia seeds were germinated in sterilized potting soil for 10 days at 21°C. Tomato seeds were surface-sterilized with 3.2% bleach for 20 min and germinated *in vitro* for 10 days at 25°C. Medicago seeds were scarified with sand paper, surface-sterilized with 3.2% bleach for 3 min and germinated *in vitro* for 5 days at 4°C and overnight at 25°C. Rice seeds were surface-sterilized with 70% ethanol for 1 min and 1% bleach for 30 min and germinated *in vitro* for 4 days at 37°C. Brachypodium seeds were surface-sterilized with 70% ethanol for 30 s and 2.5% bleach for 20 min and germinated *in vitro* for 1 week at 4°C and 3 days at 25°C. Wheat seeds were surface-sterilized in 70% ethanol for 10 min and 3.2% bleach for 20 min and germinated *in vitro* for 1 week at 4°C and 3 days at 25°C. Seedlings were transferred the into 50ml of Attapulgite Oil-Dri UK and inoculated with the following number of spores/plant of *R. irregularis* DAOM 197198 (Medicago: 200; petunia, tomato, Brachypodium and wheat: 500; rice: 1000) and watered with 0.5x modified Long Ashton medium (containing 7.5 mM NaH_2_PO_4_). For experiments including symbiotic signals, stock solutions of 10^-4^ M CO4 or LCO-V(C18:1,NMe,S) were made in water or 50% acetonitrile respectively. For treatments, 20 ml of 0.5x modified Long Ashton medium supplemented with 10^-7^ M LCO, 10^-7^ M CO4 + 0.05% acetonitrile or 0.05% acetonitrile (control) were added to the dry substrate at the beginning of the experiments and 10 ml were added twice 2 weeks and 4 weeks later. After harvest, for microscopy analysis, roots were stained in a 5% ink–vinegar solution after 10% (w/v) KOH treatment, for qRT-PCR, roots were frozen in liquid-nitrogen.

### Nodulation phenotyping

Nodule numbers were counted in individual plants grown in tubes on Fahraeus agar slants supplemented with 0.2 mM NH_4_NO_3_ at 28 dpi using 0.5 ml/tube of 0D600=0.00025 (about 1 10^5^ cfu) of *S. meliloti* strains 2011 (GMI 6526) and its *nodFE* mutant (GMI 6528), containing a constitutive pHemA:LacZ fusion.

### Calcium spiking measurement

gGECO and DsRed were respectively excited with the 488-nm or the 561-nm argon laser lines, and fluorescent emission (500 to 550 nm or 570 to 650 nm) were imaged for 20min on a SP8 confocal laser-scanning microscope (Leica) with a fluostar VISIR 25x/0.95 WATER objective. All nuclei in the focal plan were selected as regions of interest using the software LAS X and the fluorescence intensities in each region of interest were exported. Total number of cells with visible gGECO basal fluorescence and the number of cells with nuclei spiking more than once were used to calculate the percentage of cells with spiking nuclei.

### Root architecture measurement

Experiments were performed as described in^39^. Briefly, plantlets were grown in 20*20 cm plates containing Fahraeus agar medium for 2 weeks at 25°C and root systems were scanned. Total root length was estimated by using WINRhizo software and the number of lateral roots was counted manually on the images.

### LCO and CO binding assays

Western blotting, membrane fraction preparation and LCO binding assays were performed as described in ^15^. LCO-V(C18:1,NMe,S) were purified from the rhizobial strain *Rhizobium tropici*. CO4, CO5, CO7 and CO8 were prepared as described in^40^. Labelling of LCO-V(C18:1,NMe) was performed as described in ^41^. CO5-biot synthesis is described in^12^. CO5-Biot binding to membrane fractions was performed in binding buffer (25mM NaCacodylate pH 6, 1mM MgCl_2_, 1mM CaCl_2_, 250mM saccharose and protease inhibitors) for 1 hour on ice. Samples were then centrifuged for 30min at 31 000*g* and 4°C and pellets were washed and resuspended in binding buffer. Proteins were then separated by SDS PAGE, transferred to a nitrocellulose membrane, and revealed using anti-GFP antibodies (AMSBIO TP401) and after membrane stripping by streptavidin-HRP (Sigma S2438-250UG). For protein purification, membrane fractions were resuspended after centrifugation by IP buffer (25mM Tris pH7.5, 150mM NaCl, 10% glycerol, NaF protease inhibitors and phosphatase inhibitor). Proteins were then solubilized in IP buffer supplemented with 0.2% DDM for 1h at 4°C and centrifuged at 100 000*g* and 4°C for 15 min. Solubilized proteins in the supernatant were purified using *GFP-TrapMA* (ChromoTek).

### Microscale thermophoresis

Binding experiments were performed with the *Monolith NT.115* (Nano Temper Technologies). BdLYR2-ECR and BdLYR1-ECR were labelled with the *His-Tag labelling Kit RED-tris-NTA 2nd Generation* (Nano Temper Technologies) according to the manufacturer instructions, using a protein:dye ratio 2:1, in PBS buffer with Foscholine 0.175%.

For CO binding experiments, 40 nM of labelled proteins were mixed (1:1 volumes) with a range of CO serial dilutions in PBS buffer, Foscholine 0.175%, in PCR tubes, and incubated at room temperature for 10 min.

Due to the physico-chemical properties of the LCO, the above procedure was modified to avoid sticking problems and insertion into micelles. A range of LCO concentrations was prepared by serial dilution in 50% ethanol in PCR tubes. One µl of each LCO dilution was distributed in a low binding microtiter plate, 100µl of 20 nM labelled proteins were immediately added and the mix was incubated at room temperature for 10 min.

For binding experiments with Fluorescent CO5, 100nM of CO5-BODIPY (Suppl Figure 2b and 11) in MST buffer (50 mM Tris-HCl pH 7.4, 150 mM NaCl, 10 mM MgCl_2_, 0.05% tween-20) were mixed (1:1 volumes) with a range of unlabelled protein serial dilutions in MST buffer in PCR tubes, and incubated at room temperature for 10 min.

*Premium treated capillaries* (NanoTemper Technologies, for LCO and CO5-BODIPY) and *standard capillaries* (NanoTemper Technologies, for CO) were loaded and the measurements were performed at 22°C with 60 % exitation power (CO5-BODIPY) or 100% exitation power (Labelled proteins) and medium MST power. Binding data were analysed using the *MO.Affinity Analysis Software* (Nano Temper Technologies).

### qRT-PCR

Total RNA was isolated using an *NucleoSpin® RNA Plant Mini kit* (Macherey-Nagel) according to the manufacturer’s instructions. The integrity and quantity of the RNA preparations were checked using *Agilent RNA 6000 Nano Chips* (Agilent Technologies). The first-strand cDNA synthesis was performed with the *Prime Script RT Master Mix kit* (Takara). Quantitative real-time PCR was performed at 60°C using *Takyon No ROX SYBR 2X MasterMix blue dTTP* (Eurogentec) in the *LightCycler 480* (Roche) with the primers listed in Suppl data. Gene expression levels were calculated relative to the three wheat housekeeping genes listed in Suppl data. Technical duplicates were made for each biological replicate.

## Acknowledgments

We thank Fabienne Maillet (LIPME) for purification of rhizobial LCO, Sébastien Cabanac and Noémie Carles for technical help, and Sébastien Antelme for providing the Brachypodium seeds. We thank Clare Gough, Sandra Bensmihen for critical reading of the manuscript. We thank Wu Xi Shu Guang Agricultural Science and Technology Development Co. Ltd. for a PhD fellowship for Yi Ding. This work was supported by the ANR projects “NICECROPS” (ANR-14-CE18-0008-01) and “WHEATSYM” (ANR-16-CE20-0025-01), the National Natural Science Foundation of China (Grant No. 32100241) and the Foundation for Innovative Research Groups of the Natural Science Foundation of Chongqing (Grant No. cstc2021jcyj-cxttx0004). This study is set within the framework of the “Laboratoires d’Excellences (LABEX)” TULIP (ANR-10-LABX-41), SPS (ANR-10-LABX-40-SPS) and of the “École Universitaire de Recherche (EUR)” TULIP-GS (ANR-18-EURE-0019). “La Fondation pour le développement de la Chimie des substances Naturelles et ses applications” is acknowledged for granting A.M. with a PhD scolarship. S.F. acknowledges ANR for partial support through LABEX ARCANE/CBH-EUR-GS (ANR-17-EURE-0003) as well as NanoBio ICMG (UAR 2607) for providing facilities for mass spectrometry (A. Durand, L. Fort, R. Gueret). The authors declare no conflicts of interest.

## References

1. Rich, M.K., Nouri, E., Courty, P.E. & Reinhardt, D. Diet of Arbuscular Mycorrhizal Fungi: Bread and Butter? Trends Plant Sci 22, 652–660 (2017).

2. Delaux, P. et al. Comparative Phylogenomics Uncovers the Impact of Symbiotic Associations on Host Genome Evolution. Plos Genet 10, e1004487 (2014).

3. Maillet, F. et al. Fungal lipochitooligosaccharide symbiotic signals in arbuscular mycorrhiza. Nature 469, 58–63 (2011).

4. Genre, A. et al. Short-chain chitin oligomers from arbuscular mycorrhizal fungi trigger nuclear Ca2+ spiking in Medicago truncatula roots and their production is enhanced by strigolactone. New Phytol 198, 179–189 (2013).

5. Li, X.R. et al. Nutrient regulation of lipochitooligosaccharide recognition in plants via NSP1 and NSP2. Nat Commun 13, 6421 (2022).

6. Volpe, V. et al. Long-lasting impact of chitooligosaccharide application on strigolactone biosynthesis and fungal accommodation promotes arbuscular mycorrhiza in Medicago truncatula. New Phytol 237, 2316–2331 (2023).

7. Buendia, L., Girardin, A., Wang, T., Cottret, L. & Lefebvre, B. LysM Receptor-Like Kinase and LysM Receptor-Like Protein Families: An Update on Phylogeny and Functional Characterization. Front Plant Sci 9, 1531 (2018).

8. Kaku, H. et al. Plant cells recognize chitin fragments for defense signaling through a plasma membrane receptor. Proc Natl Acad Sci U S A 103, 11086–91 (2006).

9. Fliegmann, J. et al. Biochemical and phylogenetic analysis of CEBiP-like LysM domain-containing extracellular proteins in higher plants. Plant Physiology and Biochemistry 49, 709–720 (2011).

10. Cao, Y. et al. The kinase LYK5 is a major chitin receptor in Arabidopsis and forms a chitin-induced complex with related kinase CERK1. Elife 3, e03766 (2014).

11. Liu, S. et al. Molecular Mechanism for Fungal Cell Wall Recognition by Rice Chitin Receptor OsCEBiP. Structure 24, 1192–200 (2016).

12. Cullimore, J. et al. Evolution of Lipochitooligosaccharide Binding to a LysM-RLK for Nodulation in Medicago truncatula. Plant Cell Physiol 64, 746–757 (2023).

13. Bouchiba, Y. et al. An integrated approach reveals how lipo-chitooligosaccharides interact with the lysin motif receptor-like kinase MtLYR3. Protein Sci 31, e4327 (2022).

14. Broghammer, A. et al. Legume receptors perceive the rhizobial lipochitin oligosaccharide signal molecules by direct binding. Proc Natl Acad Sci U S A 109, 13859–64 (2012).

15. Girardin, A. et al. LCO Receptors Involved in Arbuscular Mycorrhiza Are Functional for Rhizobia Perception in Legumes. Curr Biol 29, 4249–4259 (2019).

16. Gysel, K. et al. Kinetic proofreading of lipochitooligosaccharides determines signal activation of symbiotic plant receptors. Proc Natl Acad Sci U S A 118, e2111031118 (2021).

17. Wang, T. et al. LysM receptor-like kinases involved in immunity perceive lipo-chitooligosaccharides in mycotrophic plants. Plant Physiol 192, 1435–1448 (2023).

18. He, J. et al. A LysM Receptor Heteromer Mediates Perception of Arbuscular Mycorrhizal Symbiotic Signal in Rice. Mol Plant 12, 1561–1576 (2019).

19. Chiu, C.H., Roszak, P., Orvošová, M. & Paszkowski, U. Arbuscular mycorrhizal fungi induce lateral root development in angiosperms via a conserved set of MAMP receptors. Curr Biol 32, 4428–4437.e3 (2022).

20. Shinya, T. et al. Functional characterization of CEBiP and CERK1 homologs in arabidopsis and rice reveals the presence of different chitin receptor systems in plants. Plant Cell Physiol 53, 1696–706 (2012).

21. Shibuya, N., Kaku, H., Kuchitsu, K. & Maliarik, M.J. Identification of a novel high-affinity binding site for N-acetylchitooligosaccharide elicitor in the membrane fraction from suspension-cultured rice cells. FEBS Lett 329, 75–8 (1993).

22. Baureithel, K., Felix, G. & Boller, T. Specific, high affinity binding of chitin fragments to tomato cells and membranes. Competitive inhibition of binding by derivatives of chitooligosaccharides and a Nod factor of Rhizobium. J Biol Chem 269, 17931–8 (1994).

23. Felix, G., Regenass, M. & Boller, T. Specific perception of subnanomolar concentrations of chitin fragments by tomato cells - induction of extracellular alkalinization, changes in protein-phosphorylation, and establishment of a refractory state. Plant Journal 4, 307–316 (1993).

24. Feng, F. et al. A combination of chitooligosaccharide and lipochitooligosaccharide recognition promotes arbuscular mycorrhizal associations in Medicago truncatula. Nat Commun 10, 5047 (2019).

25. Olah, B., Briere, C., Becard, G., Denarie, J. & Gough, C. Nod factors and a diffusible factor from arbuscular mycorrhizal fungi stimulate lateral root formation in Medicago truncatula via the DMI1/DMI2 signalling pathway. Plant Journal 44, 195–207 (2005).

26. Buendia, L. et al. Lipo-chitooligosaccharides promote lateral root formation and modify auxin homeostasis in Brachypodium distachyon. New Phytol 221, 2190–2202 (2019).

27. Ried, M.K., Antolín-Llovera, M. & Parniske, M. Spontaneous symbiotic reprogramming of plant roots triggered by receptor-like kinases. Elife 3(2014).

28. Pietraszewska-Bogiel, A. et al. Interaction of Medicago truncatula Lysin Motif Receptor-Like Kinases, NFP and LYK3, Produced in Nicotiana benthamiana Induces Defence-Like Responses. Plos One 8, e65055. (2013).

29. Huang, R. et al. Natural variation at OsCERK1 regulates arbuscular mycorrhizal symbiosis in rice. New Phytol 225, 1762–1776 (2020).

30. Vandenbussche, M. et al. Generation of a 3D indexed Petunia insertion database for reverse genetics. Plant J 54, 1105–14 (2008).

31. Dalmais, M. et al. A TILLING Platform for Functional Genomics in Brachypodium distachyon. PLoS One 8, e65503 (2013).

32. EpiTrans, I., 2018. EPIgenomics and TRANSlational Research Facility, doi: 10.15454/1.5572407597184844E12.

33. Magne, K. et al. Roles of BdUNICULME4 and BdLAXATUM-A in the non-domesticated grass Brachypodium distachyon. Plant J 103, 645–659 (2020).

34. Bendahmane, A., Marcel, F., Dalmais, M., Beaumont, G. & Mania, B. SENTINEL, SOFTWARE dedicated to TILLING by NGS Analysis. Certifier par l’Agence pour la Protection des programmes. (Inter Deposit Digital Number.FR001.240004.000.R.P.2016.000.10000).

35. Ma, X. & Liu, Y.G. CRISPR/Cas9-Based Multiplex Genome Editing in Monocot and Dicot Plants. Curr Protoc Mol Biol 115, 31.6.1–31.6.21 (2016).

36. Luu, T.B. et al. Analysis of the structure and function of the LYK cluster of Medicago truncatula A17 and R108. Plant Sci 332, 111696 (2023).

37. Cope, K.R. et al. The Ectomycorrhizal Fungus Laccaria bicolor Produces Lipochitooligosaccharides and Uses the Common Symbiosis Pathway to Colonize Populus Roots. Plant Cell 31, 2386–2410 (2019).

38. Fliegmann, J. et al. Lipo-chitooligosaccharidic Symbiotic Signals Are Recognized by LysM Receptor-Like Kinase LYR3 in the Legume Medicago truncatula. ACS Chem Biol 8, 1900–1906 (2013).

39. Bartoli, C. et al. Rhizobium leguminosarum symbiovar viciae strains are natural wheat endophytes that can stimulate root development. Environ Microbiol (2022).

40. Masselin, A. et al. Optimizing Chitin Depolymerization by Lysozyme to Long-Chain Oligosaccharides. Mar Drugs 19(2021).

41. Gressent, F., Cullimore, J.V., Ranjeva, R. & Bono, J.J. Radiolabeling of lipo-chitooligosaccharides using the NodH sulfotransferase: a two-step enzymatic procedure. BMC Biochem 5, 4 (2004).

