## Supplemental Figures for "A new group of LysM-RLKs involved in symbiotic signal perception and arbuscular mycorrhiza establishment"

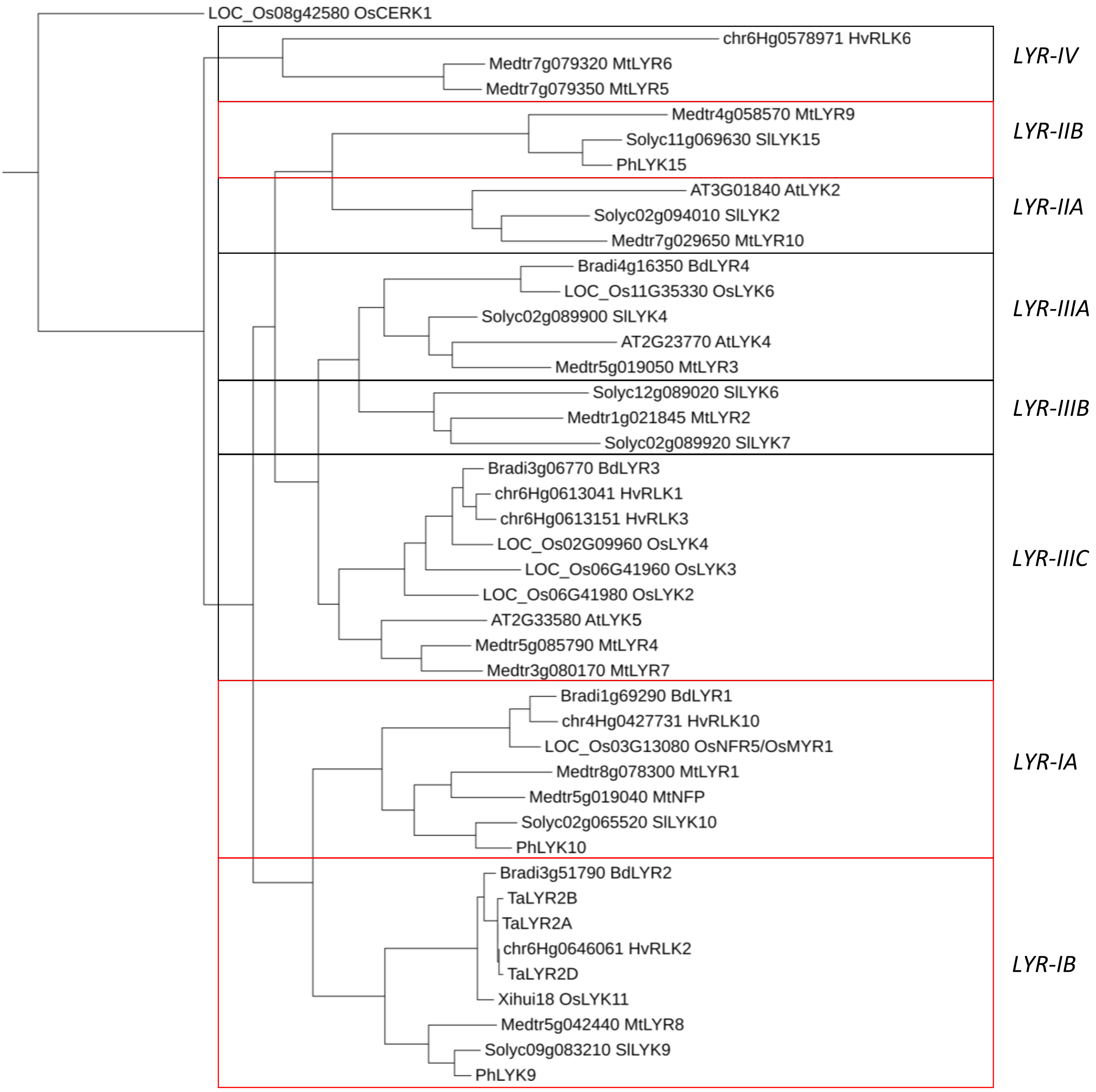


Suppl Figure 1. Phylogenetic tree of the *LYR* protein subfamily.

Complete *LYR* subfamily of *Arabidopsis thaliana*, *Medicago truncatula*, *Solanum lycopersicum Brachypodium distachyon*, *Oryza sativa* (Buendia et al., 2018) and *Hordeum vulgare* (Li et al. 2022) are shown, except SlLYK8 which is a truncated protein. *Petunia hybrida LYR-IA* and *LYR-IB* proteins as well as *Triticum aestivum* and Xihui18 *LYR-IB* proteins and the *Petunia hybrida LYR-IIB*, studied in this article, have been added. The *LYK-I*, OsCERK1, was used as an outgroup. The tree was generated with Phylogeny One click (https:://ngphylogeny fr/workflows/oneclick/) using default settings and presented using itol tree software.

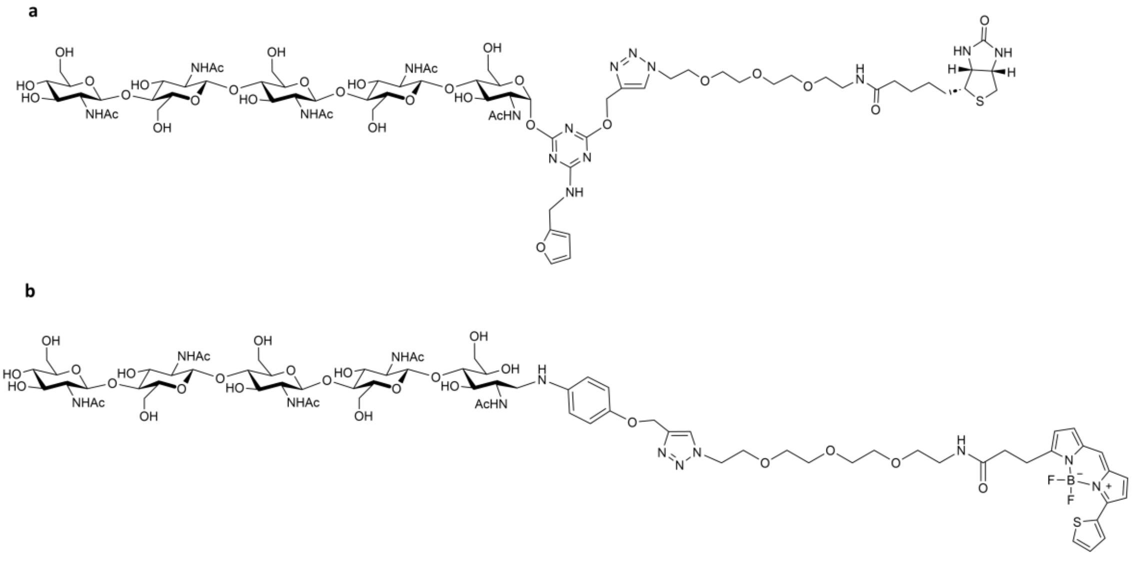


Suppl Figure 2. Structures of the modified CO used in this study.

CO5-biot (**a**) and CO5-BODIPY (**b**). The furfuryl group present on the triazine cross-linking moiety (**a**) offers the possibility to introduce an additional chemical group (fluorescent for example) on the probe but this property has not been used in this study.

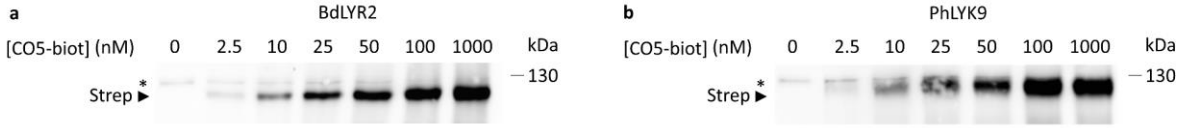


Suppl Figure 3. Affinity of PhLYK9 and BdLYR2 for CO5-biot.

Affinity of BdLYR2 (**a**) and PhLYK9 (**b**) for short-chain CO. Saturation experiments on 10 µg of membrane proteins using a range of concentrations of CO5-biot. WB only performed using streptavidin. The arrowhead indicates the position of PhLYK9-YFP or BdLYR2-YFP whereas * indicates an *N. benthamiana* endogenously biotinylated protein.

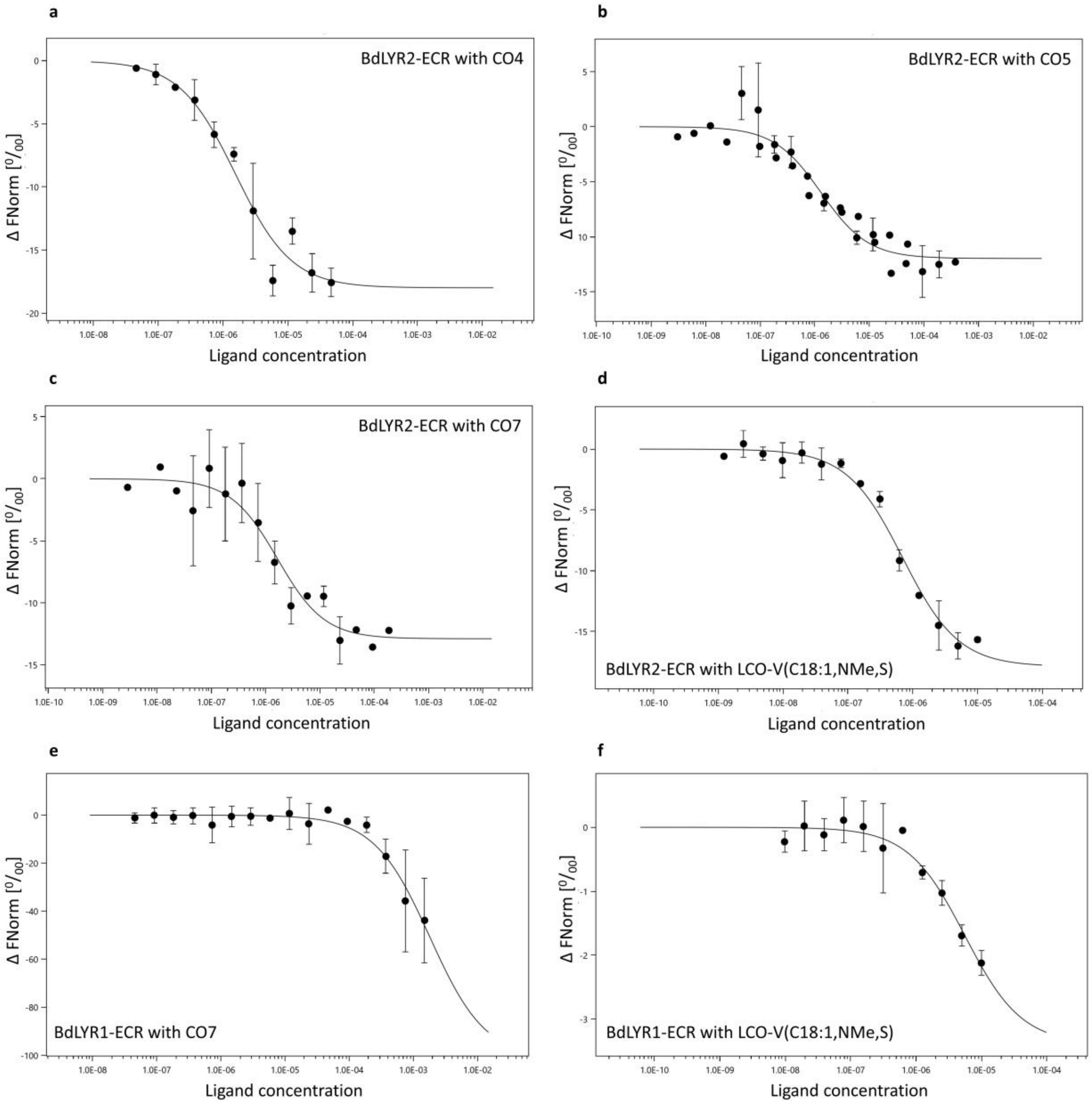


Suppl Figure 4. Microscale thermophoresis Δ FNorm plots.

Interaction between BdLYR2-ECR (**a-d**) or BdLYR1-ECR (**e-f**) and the indicated ligand. X-axes indicate ligand concentrations, and the Y-axes indicate the change in normalized fluorescence (Δ FNorm values after binding with the ligands). Error bars represent SD from two independent measurements using the same purified protein.

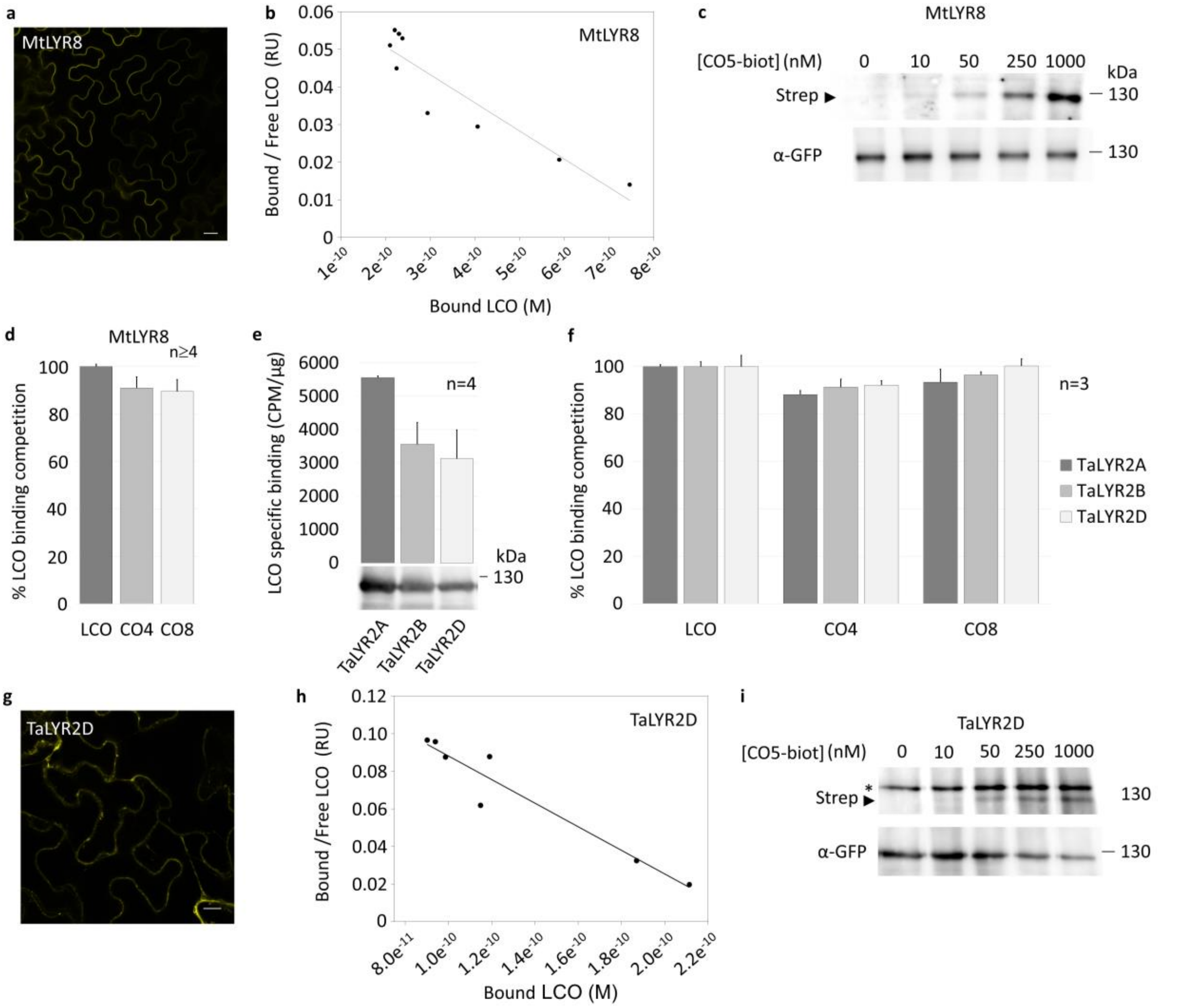


Suppl Figure 5. Affinity of MtLYR8 and TaLYR2D for LCO and short-chain CO.

**a**. Image of epidermal cells from a *N*. *benthamiana* leaf expressing MtLYR8-YFP. Scale bar represents 20 µm. **b**. Affinity of MtLYR8 for LCO-V(C18:1,NMe,S). Scatchard plot of a cold saturation experiment using membrane fractions containing MtLYR8 and a range of LCO-V(C18:1,NMe,S) concentrations as competitor. The plot is representative of experiments performed with two independent batches of membrane fractions. **c**. Affinity of MtLYR8 for short-chain CO. Saturation experiments on 200 µg of membrane proteins containing MtLYR8 using a range of concentrations of CO5-biot, followed by protein solubilization and purification using anti-GFP beads. WB performed using sequentially anti-GFP antibodies and streptavidin on the same membrane. **d**. Specificity of MtLYR8 LCO-binding sites for LCO versus CO. Bars represent the percentage competition of specific LCO-V(C18:1,NMe,$^{35}$S) binding (means and SD between at least 2 technical replicates on 2 batches of membrane fractions) in the presence of 2 µM of LCO-V(C18:1,NMe,S), CO4 or CO8. **e**. Specific binding of LCO-V(C18:1,NMe,$^{35}$S) to TaLYR2A, TaLYR2B and TaLYR2D. Bars represent the specific LCO-binding/µg membrane proteins (means and SD from 4 technical replicates on 1 batch of membrane fractions containing the indicated proteins). Immunodetection with anti YFP antibodies in 10µg of the indicated membrane fractions. **f**. Specificity of TaLYR2A, TaLYR2B and TaLYR2D LCO-binding sites for LCO versus CO as in **e**. **g**. Image of epidermal cells from a *N*. *benthamiana* leaf expressing TaLYR2D-YFP as in **a**. **h**. Affinity of TaLYR2D for LCO-V(C18:1,NMe,S) as in **b**. **i**. Affinity of TaLYR2D for short-chain CO. Saturation experiments on 10 µg of membrane proteins containing MtLYR8 using a range of concentrations of CO5-biot. WB as in **c.** The arrowhead indicates the position of TaLYR2D-YFP whereas * indicates an *N. benthamiana* endogenously biotinylated protein.

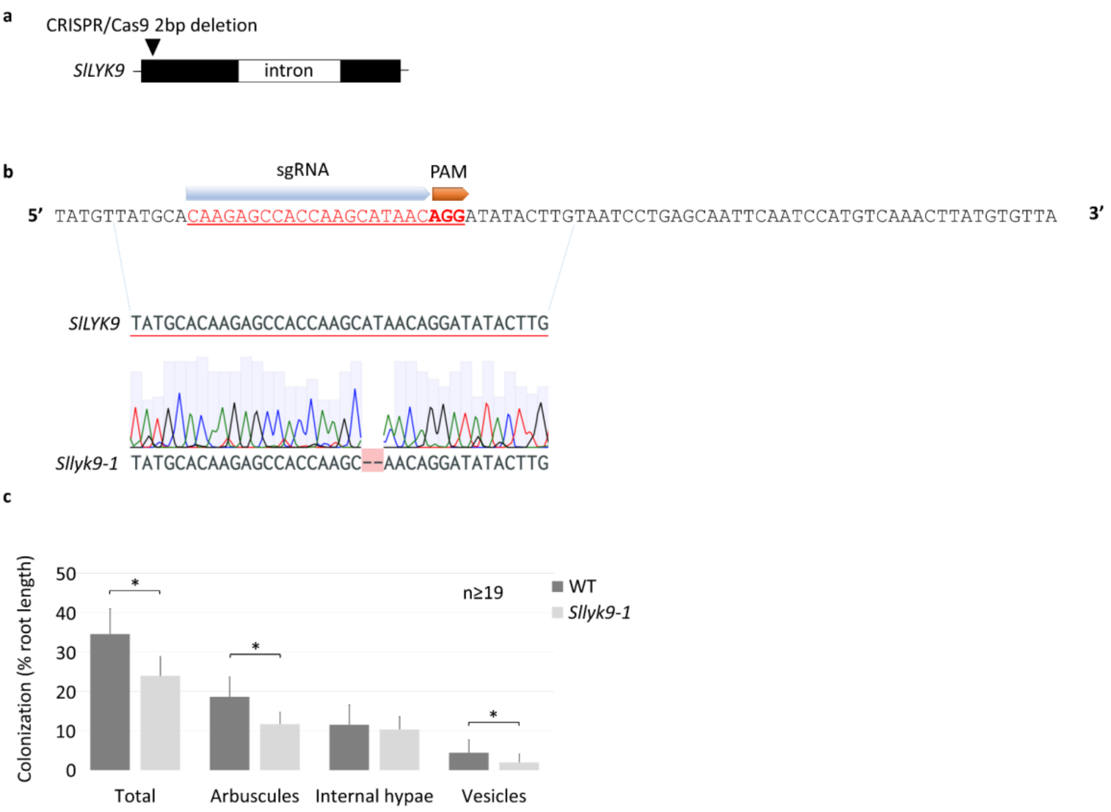


Suppl Figure 6. CRISPR-CAS9 edition and AM phenotype of tomato *Sllyk9-1*.

**a.** Position of the frameshift deletion in *Sllyk9-1* in the Ailsa Craig cultivar. **b.** Representative trace of T2 plant sequencing. **c.** Detailed analysis of AMF structures at 4 wpi in *Sllyk9-1* and WT Ailsa Craig plants. Means and SD between root systems from 2 independent experiments are shown. Statistical differences (p-value < 0.05) were calculated using pairwise a T test.

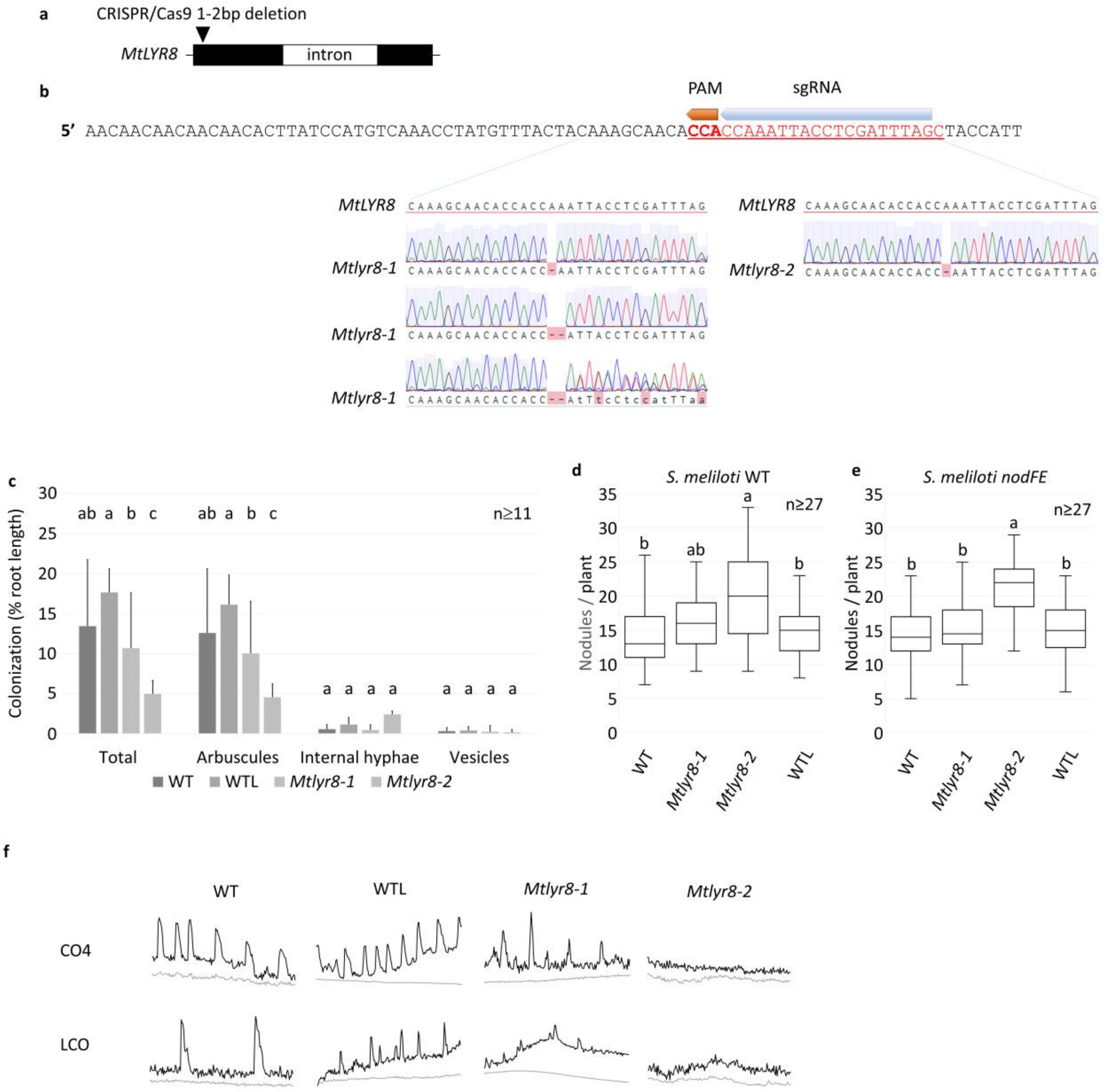


Suppl Figure 7. CRISPR-CAS9 edition and symbiosis phenotypes of *Mtlyr8-1* and *Mtlyr8-2*.

**a.** Position of the frameshift deletions in *Mtlyr8-1* and *Mtlyr8-2* in the 2HA accession. **b.** Protospacer sequence and representative traces of individual T2 plant sequencing. *Mtlyr8-1* is a bi-allelic segregating mutant (deletion of 1 and 2 bp), whereas *Mtlyr8-2* has a 1 bp deletion. **c.** Detailed analysis of AMF structures at 4 wpi in *Mtlyr8-1*, *Mtlyr8-2*, *A17* (WT) or a 2HA line transformed with an empty vector (WTL). Means and SD between root systems from 1 experiment are shown. **d-e.** The number of nodules at 28 dpi with WT (d) or *nodFE* (e) *S. meliloti* in *Mtlyr8-1*, *Mtlyr8-2*, 2HA (WT) or a 2HA line transformed with an empty vector (WTL). Boxplots represent the distribution between root systems from 2 independent experiments. Statistical differences (p-value < 0.05) were calculated using a van der Waerden test. **f.** Representative green (black line) and red (grey line) fluorescence traces over time (x-axis, 20 min) in nuclei of transgenic G-GECO-DsRed roots from the indicated lines following treatment with $10^{-7}$ M CO4 or $10^{-7}$ M LCO-V(C18:1,NMe,S).

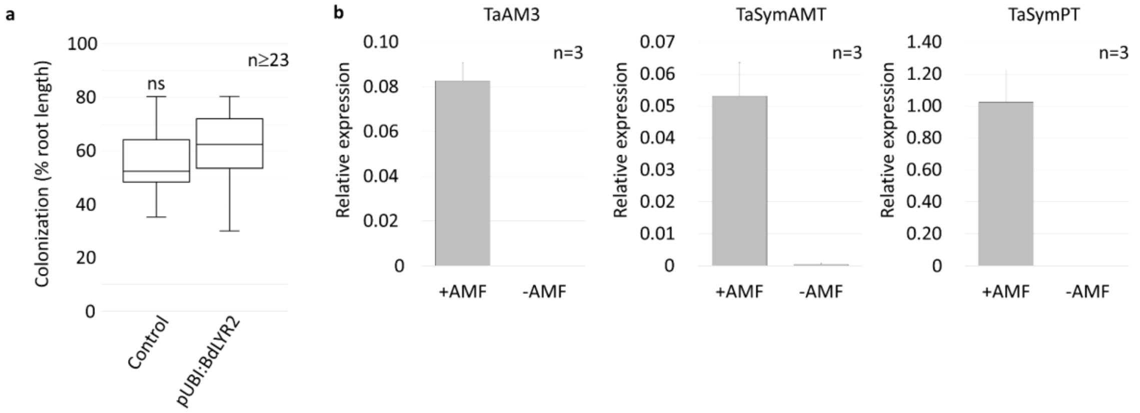


Suppl Figure 8. AMF colonization in wheat expressing BdLYR2 and validation of wheat AM marker genes

**a.** Root-length colonization at 46 dpi in wheat plants expressing BdLYR2 under the control of a strong promoter (pUbi:BdLYR2) or containing the empty vector (Control). Boxplots represent the distribution between root systems of 2 independent transgenic lines for each construct. No significant differences (ns) were found between using a T test (p-value < 0.05). **b.** Relative expression of the early AM stage marker genes TaAM3 and the ammonium and phosphate transporters specifically expressed in arbuscule-containing cells (SymAMT2 and symPT respectively) in a wheat cultivar (Renan) inoculated or not with AMF at 4 wpi. Boxplots represent the distribution between 3 pools of roots.

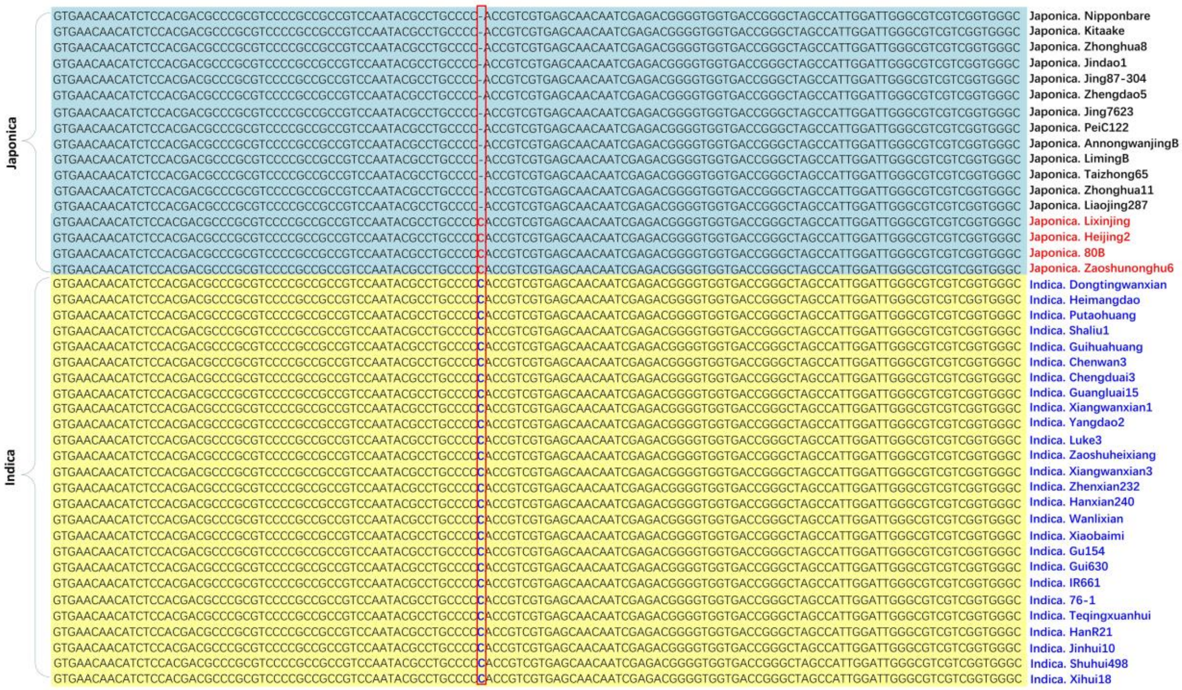

Suppl Figure 9. Presence/absence of the deletion in *OsLYK11* in various rice indica and japonica cultivars.

Alignment of *OsLYK11* sequences surrounding the position of the deletion (boxed in red), creating a frame shift in some rice cultivars. Sequences highlighted in yellow are from indica cultivars, those in turquoise from japonica cultivars. None of the indica cultivars has the deletion. Most of the japonica cultivars have the deletion (cultivar names in black), but some of them do not (cultivar names in red).

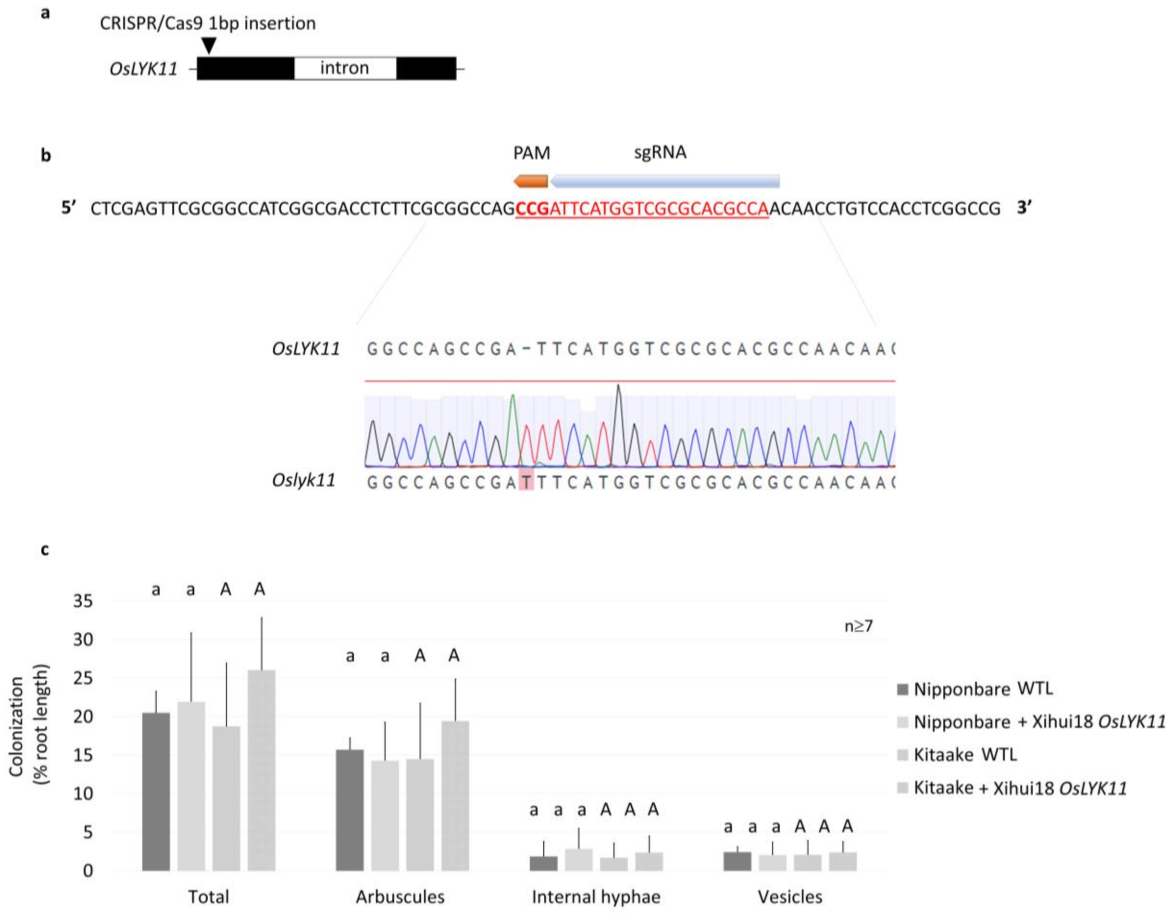


Suppl Figure 10. CRISPR-CAS9 edition and AM phenotype of *Oslyk11-1*.

**a.** Position of the frameshift insertion in *Oslyk11-1* in the Xihui18 cultivar. **b.** Representative trace of T2 plant sequencing. **c.** Detailed analysis of AMF structures at 5 wpi in Nipponbare or Kitaake plants containing the empty vector (WTL) or Xihui18 *OsLYK11*. Means and SD between roots systems of 2 independent transgenic lines from 1 experiment. Statistical differences (p-value < 0.05) were calculated using a van der Waerden test.

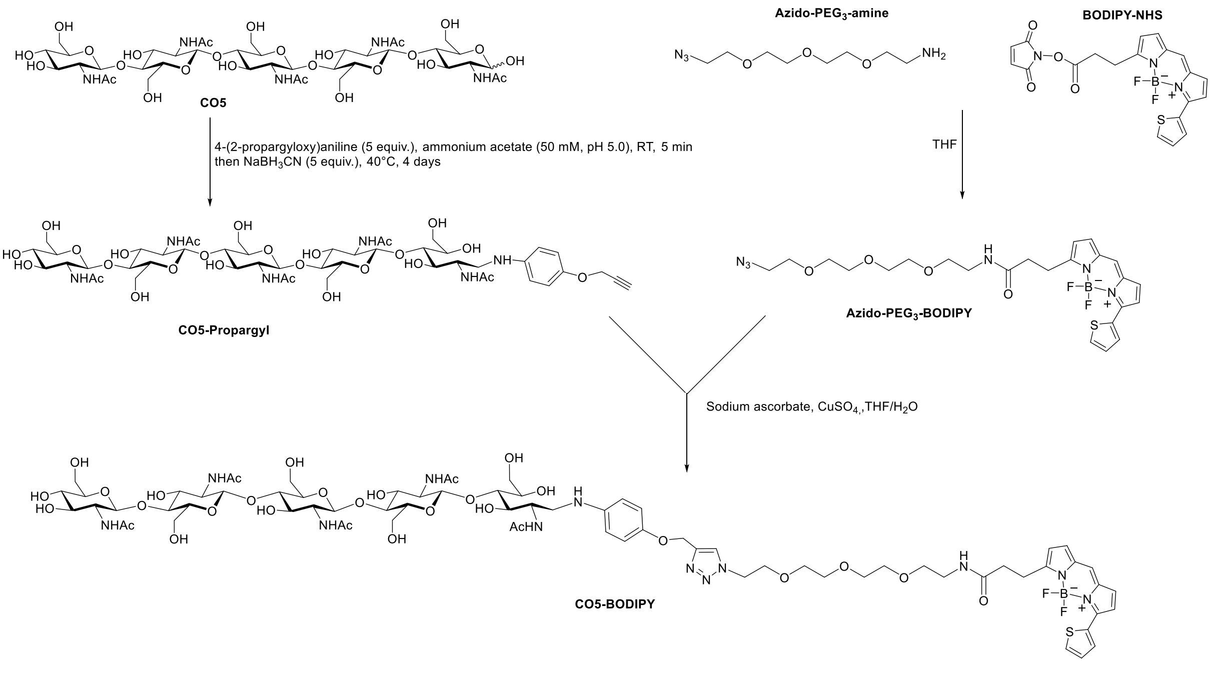


Suppl Figure 11. CO5-BODIPY synthesis.

BODIPY-NHS (BDP 558/568 NHS ester, CAS 150173-73-2) was purchased from Lumiprobe (Hannover, Germany). 4-(2-Propynyloxy)aniline (CAS 26557-78-8) and Azido-PEG$_3$-amine (CAS 134179-38-7) were purchased from TCI Europe (Zwijndrecht, Belgium).

**CO5-propargyl**

The reaction conditions were previously reported by Guerry et al[1]. 4-(2-Propynyloxy)aniline (39 mg, 0.27 mmol) in MeOH (300 µL) was added to a stirred solution of CO5 (56 mg, 54 µmol) in 50 mM ammonium acetate buffer pH 5.0 (1.7 mL). The reaction was stirred at room temperature for 5 min and $NaBH_3CN$ (17 mg, 0.27 mmol) was added. The reaction mixture was then stirred at 40°C during 4 days. The solution was concentrated and the residue was purified by C18 reversed-phase chromatography on a SPE cartridge. CO5-propargyl was isolated with a yield of 57% (36 mg). HRMS (ESI) *m/z* calcd for $C_{49}H_{76}O_{26}N_6$+$2H^+$: 583.24774 $[M+2H]^{2+}$; found: 583.24673.

**Azido-PEG$_3$-BODIPY**

Azido-PEG$_3$-amine (5 µL, 25 µmol) in water (50 µL) was added to BODIPY-NHS (10 mg, 23 µmol) in THF (1 mL). The reaction was stirred overnight at room temperature and then concentrated under vacuum. The residue was taken up in ethyl acetate (5 mL) and successively washed with water (5 mL), 1N HCl (5 mL) and 1N NaOH (5 mL). The organic phase was dried over anhydrous sodium sulfate, filtrated and concentrated. Azido-PEG$_3$-BODIPY was isolated with a yield of 80% (10 mg), characterized by mass spectrometry (*m/z* : 569 $[M+Na]^+$) and directly used for the next step without further purification.

**CO5-BODIPY**

Azido-PEG$_3$-BODIPY (3.5 mg, 6.4 µmol) in THF (350 µL) was mixed to a solution of CO5-propargyl (8 mg, 7 µmol) in water (400 µL). Aqueous solutions of sodium ascorbate (10 mg/mL, 90 µL, 4.5 µmol) and copper sulfate (10 mg/mL, 70 µL, 4.5 µmol) were successively added and the reaction mixture was stirred overnight at room temperature. The solution was concentrated and the residue was purified by C18 reversed-phase chromatography on a SPE cartridge. CO5-BODIPY was isolated with a yield of 28% (3 mg). HRMS (ESI) *m/z* calcd for $C_{73}H_{105}O_{30}N_{12}BF_2S$+$2H^+$: 856.34994 $[M+2H]^{2+}$; found: 856.34978.

1. Guerry, A., Bernard, J., Samain, E., Fleury, E., Cottaz, S. & Halila, S. *Bioconjug. Chem.* **24**, 544-549 (2013).
